# Enhancing Plant Immune Training and Protection through Damage- and Microbe-Associated Molecular Patterns from Anaerobic Digestate

**DOI:** 10.1101/2025.03.03.638737

**Authors:** Marco Greco, Daniele Coculo, Angela Conti, Marco Abatematteo, Savino Agresti, Daniela Pontiggia, Hugo Mélida, Lorenzo Favaro, Vincenzo Lionetti

**Author notes:** Corresponding author: Vincenzo Lionetti.

## Abstract

Olive oil production is a major global agricultural industry that generates significant waste, particularly olive pomace, which poses environmental and economic challenges. Anaerobic digestion has emerged as a promising solution for its valorization into biogas and reducing its environmental impact. However, the resulting digestate remains underutilized and its long-term environmental impact is uncertain. Traditional disposal methods are costly and inefficient, underscoring the need for more sustainable approaches. In this study, olive pomace digestate was biorefined and its components were upcycled into soil amendments and plant immunostimulants. Metagenomic analysis revealed a diverse microbial community in the liquid fraction, including *Luteimonas*, *Pseudomonas*, and *Caldicoprobacter*. We obtained a MIcrobial Protein Extract (MIPE) from this biomass, containing precursors of microbe- and damage-associated molecular patterns including Flagellin, Elongation Factor Tu, and the phytocytokine Golven. Treatment with MIPE triggered a rapid plant immune response, characterized by increased hydrogen peroxide production, phosphorylation of mitogen-activated protein kinases, and the upregulation of defense-related genes such as *CYP81F2*, *FRK1*, and *WRKY53.* MIPE-induced priming enhanced Arabidopsis and tomato resistance to *Botrytis cinerea* and *Pseudomonas syringae*. Our findings highlight digestate as a source of bioelicitors, offering a sustainable alternative to chemical pesticides while enhancing plant immunity, valorizing olive mill waste and promoting sustainable agriculture.

## 1. INTRODUCTION

A significant worldwide rise in the extraction and consumption of olive oil has been recorded, driven by its appealing organoleptic qualities and the growing acknowledgment of its health benefits ^1^. However, managing olive mill wastes (OMW), including pomace, stones, and leaves, is quite challenging, with current methods such as composting and specialized treatment involving high costs ^2^. In particular, inadequate management of two-phase pomace, an acidic liquid rich in organic compounds, polyphenols, and heavy metals, leads to improper disposal, exacerbating soil degradation and water eutrophication ^3^. Developing sustainable methods to convert OMW into valuable bioproducts is crucial to reduce costs and minimize environmental impact ^4,5^. Anaerobic digestion (AD) is gaining attention as a promising biological solution for valorizing OMW by converting it into biogas and bioenergy ^6,7^. This process not only mitigates greenhouse gas emissions but also enhances the profitability of the olive oil industry ^8–10^. As a result, more olive oil mills are adopting biogas plants to leverage olive pomace as a valuable biomass resource in anaerobic digestion systems ^11,12^. AD can convert complex organic matter into biogas through stages such as hydrolysis, acidogenesis, acetogenesis, and methanogenesis, each facilitated by a diverse microbial community of bacteria and archaea working in coordination ^13,14^. While extensive research has focused on optimizing anaerobic digestion conditions, comparatively less attention has been given to managing the digestate, the semi-solid byproduct of the AD process.

Nowadays, digestate disposal represents a key challenge for the biogas industry. Digestate remains an industrial byproduct that is underutilized and inadequately managed. Although digestate can be dried and incinerated, this method emits harmful carbon gases into the atmosphere. A notable limitation of digestate is its high moisture content, which complicates the transportation and distribution of its nutrient constituents. The absence of effective strategies for sustainable digestate management may severely impede the development of the biogas industry ^15,16^. A potential benefit comes from using digestate as a fertilizer. Digestate primarily comprises partially degraded organic components, water, and essential nutrients. Its solid fraction is enriched with recalcitrant plant organic matter. The liquid digestate (LD) predominantly contains inorganic constituents such as nitrogen, phosphorus, and potassium, in addition to various micronutrients and the microbial biomass involved in the anaerobic digestion process ^17,18^. However, the nutrient composition of digestate may be unbalanced when used as a fertilizer. Moreover, amending the soil with digestate may lead to environmental issues, notably the emission of ammonia and greenhouse gases, nutrient leaching, and the potential dissemination of pathogens that may have proliferated during the production process, along with the presence of micro-pollutants ^15,19^. Digestate can harbour a diverse community of agronomically beneficial microorganisms, including plant growth-promoting bacteria, denitrifying and nitrifying bacteria, and nitrogen-fixing bacteria ^20^. Arbuscular mycorrhizal fungi and saprophytic fungi can also be present in digestate due to their resilience as spores or contamination from raw materials and the environment ^21^. However, the digestate may also contain pathogenic bacteria, opportunistic fungi, and antibiotic-resistant microbes, which could pose risks to plant health, soil quality, and even human and animal safety if not properly managed ^22^.

However, the digestate may also contain pathogenic bacteria, opportunistic fungi, and antibiotic-resistant microbes, which could pose risks to plant health, soil quality, and even human and animal safety if not properly managed. One possible solution could be the conversion of such a biodiversity into a source of elicitors, useful in boosting plant immunity. In fact, another major challenge affecting agriculture, including olive cultivation, is the widespread use of chemical phytosanitary products. Farmers depend heavily on chemical pesticides to prevent substantial crop losses from plant pathogens ^23^. However, pesticides can pose risks to human health, particularly for agricultural workers exposed during farm activities, and for consumers due to potential pesticide residues in food ^24,25^. The plant immune system detects pathogenic microbes through pattern-recognition receptors (PRRs), which are either receptor kinases (RKs) or receptor proteins (RPs) located on the cell surface ^26,27^. These PRRs recognize conserved molecules from microbes, nematodes, insects, and parasitic plants, known as microbe-associated molecular patterns (MAMPs), triggering Pattern-Triggered Immunity (PTI) ^28,29^. MAMPs include fragments of flagellin, translation elongation factor EF-Tu, β-glucans, chitin, ergosterol, lipopolysaccharide, elicitin, and hairpin ^30–35^. Importantly, danger signals can also arise from immunogenic plant host factors ^36^. PRRs can also detect endogenous danger molecules, including phytocytokines, cytosolic proteins, peptides, nucleotides, amino acids, and Damage Associated Molecular Patterns (DAMPs), including oligosaccharides, from the degradation of the plant cell wall ^5,37,38^. Pectin fragments, named Oligogalacturonides (OG) can be released during infection and act as elicitors ^39^. MAMPs and DAMPs application to plants can confer a greater ability to detect pathogens and activate defence responses faster than untreated crops ^40^. This is a consequence of priming, a sensory state induced by the elicitors, remembering past elicitation events, and responding to the next stresses with faster and stronger activation of defence ^41,42^. However, excessive applications of these molecules can induce a hyperimmune response, causing the plant to strike a growth-defence trade-off ^5,43^.

In this study, we propose a biorefining approach to separate digestate into its components and investigate the upcycling the liquid digestate (LD) and solid digestate (SD) fractions individually for use as soil amendments, plant fertilizers, and immunostimulants. A detailed characterization of the biomass and nutrients was performed and the ability to stimulate the growth and productivity of Arabidopsis and tomato plants was compared. We explored digestate as a continuously proliferating reservoir of low-cost MAMPs/DAMPs-based plant elicitors. We isolated the microbial and plant residual biomasses from the liquid fraction of digestate produced by a digester exclusively fed with olive pomace biomass. The microbial fraction was characterized using an advanced DNA metabarcoding approach, employing 16S and ITS rRNA sequencing to comprehensively assess the diversity and composition of the microorganisms occurring in the liquid digestate. A fraction enriched in proteins/peptides, henceforth referred to as MIcrobial Protein Extract (MIPE) was obtained. The identified microbial taxa were used as a custom database for a targeted proteomic analysis to identify proteins enriched in the preparation. The preparation was tested on Arabidopsis and tomato plants to assess its potential to induce defence responses and elicit protection against pathogens such as *Botrytis cinerea* and *Pseudomonas syringae*.

## **2** Materials and methods

### 2.1. Olive pomace digestate collection

The digestate was collected from the <u>AGROLIO-AGROENERGY</u> biogas plant (Andria, BT, Italy), the first AD plant in the world to use OMW as sole feedstock. The raw digestate was generated under mesophilic conditions (45 °C) after 60 days of AD of olive pomace, derived exclusively from a two-phase olive oil extraction system processing olive (*Olea europaea* L., cultivar Coratina).

### 2.2 DNA extraction and 16S and ITS rRNA sequencing

The genomic DNA of the microbial communities in MIPE was extracted with the DNeasy PowerSoil kit (QIAGEN, Germany). In brief, the microbial pellet obtained as above mentioned was resuspended in 500 µL UREA buffer (8 M urea, 0.5 M NaCl, 20 mM Tris, 20 mM EDTA) and incubated for 30 min at 60°C, then centrifuged for 5 minutes at 10000*xg*. The supernatant was discarded, and the pellet was resuspended with 800 µL of CD1 (© QIAGEN) provided with Kit. DNA extraction followed the manufacturer’s instructions for the kit mentioned above. DNA concentration was assessed with NanoDrop1000 (ThermoFisher Scientific, MA, USA). Metagenomic DNA was used as a template for PCR amplification of the standard barcode regions currently employed in microbial taxonomy. The whole 16S was amplified with the primers pair 8F (5’-AGAGTTTGATCCTGGCTCAG) ^44^ and 1492R (5’-GGTTACCTTGTTACGACTT) ^45^. Platinum™ SuperFi II PCR Master Mix (Invitrogen™) was used for the amplification that was carried out as follows: initial denaturation at 98 °C for 30 sec, 30 amplification cycles (98 °C for 30 sec., 60 °C for 1 min, and 72 °C for 45 sec) and a final extension at 72 °C for 5 min. Amplicons were checked on 1% Agarose gel. Amplicons were then subjected to a tagging step with modified primers tailed as indicated by Oxford nanopore protocol (SQK-LSK114). Amplification was performed with Platinum™ SuperFi II PCR Master Mix (Invitrogen™) at the following conditions: initial denaturation at 98 °C for 30 sec, 15 amplification cycles (98 ° C for 10 sec., 60° C for 10 sec min and 72 °C for 45 sec) and a final extension at 72 °C for 5 min. Amplicons were checked on 1% Agarose gel. PCR products were size-selected by cleaning up with 0.7× volume of Ampure XP (Beckman Coulter, Brea, CA, USA). 200 fmol of each sample was used for the barcoding step, according to the Ligation sequencing Kit (SQK-LSK114) and the PCR Barcoding Expansion Pack 1-96 (EXP-PBC096) protocol (Oxford Nanopore Technologies, Oxford, UK). After a purification step with 0.7X Ampure XP, a pooled barcoded library was prepared by mixing 10.46 ng of DNA per sample to reach the final concentration of 1 µg of DNA in 47 µL of nuclease-free water. The library was end-repaired and adapted for nanopore sequencing by using the NEBNext Ultra DNA library preparation kit. 50fmol of the product was loaded onto the R10.4.1 flow cell. The quantification steps were carried out in NanoDrop TM 1000 (ThermoFisher Scientific, MA, USA). FASTA5 produced with MinION was called Guppy (version 6.4.6) on a supported NVIDIA GPU. The minimum quality threshold of reads to be considered for the analysis was set to 10. Passed raw reads were filtered using seqtk to remove sequences shorter than 400 bp and longer than 1800 bp. Filtered reads were merged into a single file, which served as the input for the alignment program minimap2 (version 2.24), aligning against SILVA 16S database (SILVA_138.1_SSURef_tax_silva.fasta.gz). The mapping algorithm was tuned to support the alignment of long-noisy reads using the option map-ont, which uses ordinary minimizers as seeds. Results were stored in SAM files processed with the SAMtools package to get tab-delimited tables. The final tables consisted of a list of reference strains with an indication of the number of raw reads that were assigned (mapped) to each strain. To account for variable library sizes, data were normalized using the Geometric Mean of Pairwise Ratios (GMPR) available with the function “trans_norm” within the R package microeco, which ensures a robust normalization method for zero-inflated count data. The taxonomic abundance was calculated with the function “trans_abund” (microeco, R). Package ggplot2 was used to create bar plot and donut charts. Plots showed the compositions of the community at the Genus level and represented the mean relative abundance across all the ten replicates subjected to the metabarcoding analysis. These values were expressed with the option “use_percentage” set to TRUE. Seeing the complexity of the matrix, a considerable number of reads were identified as belonging to “uncultured” genera. To give an accurate description of the community, the abundance matrix was restricted to those genera labeled as “uncultured” and a composition analysis at Family, Order, and Phylum level was carried out. Taxonomic abundance transformations and visualization were performed with the function “trans_abund” (microeco, R). 16S and Internal Transcribed Spacer (ITS) ribosomal RNA sequences are stored in the SRA archive with the BioProject ID PRJNA1211516.

### 2.3 MIPE extraction and quantification

The protein/peptide fraction MIPE was obtained through sedimentation, centrifugation, and sonication steps, as outlined in Patent Application No. WO2024188899A1 ^46^. The presence of microbial cells was confirmed through staining by fluorescein isothiocyanate and observation by fluorescence microscopy using the Nikon Eclipse E200 microscope at 40x magnification. Images were taken with a Nikon Digital Sight DS-Fi1c camera. The microbial pellet was then resuspended in sterile distilled H₂O and total proteins were extracted by cell lysis using a sonicator (Sonics Vibra Cell VCX 130, Sonics & Materials, Inc., Newtown, CT, United States). Sonication was performed at 70% amplitude for 5 min in an ice bath to prevent overheating, using 20-second ON and 20-second OFF pulses to ensure effective disruption of microbial cells while minimizing sample degradation. The lysis mixture was then centrifuged at 15000 *x g* for 30 min at 4°C to remove microbial and plant residues and the supernatant was recovered. The supernatant was treated with 20% v/v trichloroacetic acid (Sigma Chemical Co, St. Louis, MO) in ice for 30 min to precipitate proteins. The precipitate was pelleted down by centrifugation at 6000 *x g* for 10 min at 4 °C. The precipitated protein was washed with 1.25 M HCl in ethanol to remove traces of TCA ^47^. Precipitate proteins were dried using a lyophilizer and stored at - 20 °C for subsequent analyses. Total protein concentration was estimated using the Bradford method ^48^ using a Bio-Rad protein assay kit. (Bio-Rad Laboratories Inc., Hercules, CA, USA), and bovine serum albumin was used as a standard. The precipitated proteins were analyzed by SDS polyacrylamide gel electrophoresis (SDS-PAGE) using 12% polyacrylamide gels under reducing conditions and electrophoresed at 80 V for 10 min followed by 140 V for 60 min. Proteins were visualized by Comassie brilliant blue staining. MIPE yields were quantified as the ratio between MIPE dry weight (mg) and the volume of supernatant liquid digestate after sedimentation (L). The presence of nucleic acid was evaluated by measurement of the absorbance ratios 260/280 and 260/230 nm using Thermo Scientific NanoDrop Spectrophotometer. The presence of nucleic acids was explored by electrophoresis on a 1.5% (w/v) agarose gel carried out at 60 V for 25 min. Images were captured using Gel Doc™ XR + System (BioRad, Hercules, CA, USA).

### 2.4 Protein identification by LC-MS/MS analysis

For protein identification analysis, 15 µg total proteins for 3 biological replicates of MIPE were incubated in LaemmLi loading buffer at 100 °C, then loaded onto a 12% acrylamide gel for separation by 1D-SDS-PAGE. Proteins were visualized by Coomassie Brilliant Blue staining. Each lane was cut into 7 different gel slices for in-gel trypsin digestion, performed as previously described ^49^. As previously described, peptides were analyzed by ultra-high-performance liquid chromatography coupled with high-resolution mass spectrometry ^50^. Specifically, peptides were separated using a 75 μm C18 column (ES800-PepMap™ RSLC C18, 150 mm × 75 μm, Thermo Fisher Scientific) and nano-liquid chromatography (UltiMate 3000 RSLC nano-LC system, Thermo Fisher Scientific) with 100-minute gradient elution (from 4% to 90% eluent B, 80% ACN, 0.1% formic acid, at a constant flow rate of 0.3 μL/min). They were analyzed using a Q Exactive Plus™ Hybrid Quadrupole-Orbitrap™ mass spectrometer (Thermo Fisher Scientific) under data-dependent acquisition, selecting the 15 most intense ions. Full-scan spectra ranged from m/z 350.0 to 1700.0, with a resolution of 70,000 ppm and HCD fragmentation was performed at 17,500 ppm. Protein identification was performed in the MaxQuant software v. 2.2.0.0 ^51^ using the following parameters: protein and peptide false discovery rate (FDR) < 0.01, posterior error probability based on Mascot score, minimum peptide length of 7, as previously described ^52^. A custom database (protein counts: 157496) was built for protein identification, combining the UniProtKB reference proteomes released in data 09/05/2024 for bacteria, fungi, and olive. The mass spectrometry proteomics data have been deposited to the ProteomeXchange Consortium via the jPOST partner repository [http://repository.jpostdb.org] with the dataset identifier JPST003545/PXD059822 ^53^.

### 2.5 Plant growth conditions

*A. thaliana* wild-type Columbia (Col-0) seeds were washed in 2 mL of isopropanol for 30 sec and washed in 2 mL of ultrapure sterile water for 3 min in slow agitation. Seeds were sterilized with 2 mL of sterilization solution (20% NaClO in ultrapure sterile water) for 5 min in slow agitation, followed by 4 washing steps in 2 mL of ultrapure sterile water. Seeds were stored in the dark at 4°C for 2 days to promote germination. For seedling treatments, seeds were germinated in multiwell plates (approximately 10 seeds/well) containing 1 mL per well of liquid Murashige and Skoog (MS)/2 medium Sigma (2.2 g/l MS Medium basal salt, 0.5% sucrose, pH 5.7) ^54^. To evaluate the dose effect of MIPE on Arabidopsis seedling growth, 7-day-old Arabidopsis seedlings were grown in MS/2 liquid medium containing sterile water or MIPE at different concentrations (1, 10, or 100μg/mL). Extracts were sterilized by vacuum filtering through an MCE membrane filter (0.22 µm pore size). For adult plant treatments, Arabidopsis wild-type Col-0 seeds were grown in solid MS/2 medium (2.2 g L^-1^ MS Medium basal salt, 1% sucrose, 0.8% plant agar, pH 5.7). Both plates were incubated in a controlled environmental chamber maintained at 22°C with a 16-h light/8-h dark cycle and a light intensity of 120 mmol m^-2^ s^-1^. 7-day-old seedlings grown in solid MS/2 medium were transferred in sterile soil in a growth chamber at 22°C with a 12 h light/12 h dark cycle (PAR level of 100 μmol m^-2^ s^-1^). Tomato (*Solanum lycopersicum*, Minibel) seeds were germinated on wet paper overnight, transferred on soil, and grown in the greenhouse under 16 h light/8 h dark cycle (PAR level of 75 μmol m^-2^s^-1^) at 23°C and 35-40% humidity.

### 2.6 Digestate characterization

The two-phase olive pomace raw digestate was subjected to sedimentation overnight at 4°C to separate it into liquid and solid digestate fractions. Both fractions were collected, and alcohol-insoluble residues were extracted. The monosaccharide composition of these residues was analyzed following the previously described method ^55^. The percentage of organic matter on solid fraction was evaluated by standard protocols defined by D.M. 25/03/2002, G.U. n°84, 10/04/2002. The pH of the liquid digestate was determined by using y using a CRISON GLP21 pH-meter (Hach Lange Spain, S.L.U., Barcelona, Spain). The content of nitrogen, ammonium, and ammonium nitrogen was evaluated according to the Italian standard methods ^56^. The content of potassium, phosphorus, sulfur, magnesium, iron, copper, boron, manganese, nickel, chromium hexavalent, lead, cadmium, mercury and zinc were measured according to the UNI EN 16174:2012 and UNI EN 16170:2016 methods. For oligosaccharides characterization, liquid digestate was treated as previously reported ^5^. The oligosaccharide profile of LDE was investigated by HPAEC-PAD, as previously reported ^57^.

### 2.7 Plant growth experiment

A. *thaliana* was grown in soil amended with 45, 90, or 180 g/kg of olive pomace raw digestate and after two weeks, the rosette fresh weights were measured and compared with watered plants used as control. Next, Arabidopsis was grown in soil amended with 90 g/kg of olive pomace raw digestate or with amounts of solid (30 g/kg) or liquid (60 g/kg) fractions corresponding to the relative proportions of these components present in the raw digestate. After two weeks, the rosette fresh weights were measured and compared with watered plants used as mock. *Arabidopsis* plants were grown in soil amended with liquid digestate, microbe-depleted liquid digestate (both 30 g/kg), or microbial pellet (1.5 g/kg) where the amounts of Microbial-enriched pellet (M) corresponded to the relative proportions present in the LD. After two weeks, the rosette fresh weights were measured and compared with watered plants used as mock. Five-week-old tomato plants (Minibel) were transplanted in soil amended with 20, 40, or 80 kg/m^2^ of raw olive pomace digestate and after four months, tomato height was measured and compared with untreated plants used as control. Next, tomato plants were transplanted in soil amended with 20 kg/m^2^ of olive pomace raw digestate or with amounts of solid (6 kg/m^2^) or liquid (14 kg/m^2^) fractions. The tomato height and fruit number were measured four months after the transplantation and compared with untreated plants used as control. Ten biological replicates (considering an individual plant as one biological replicate) were used for each experiment. The experiments were performed three times with similar results.

### 2.8 Fungi and bacteria cultivation

The strain *B*. *cinerea* SF1 ^58^ was grown in the dark before conidium collection at 23 °C and 70% relative humidity for 20 days on Malt Extract Agar (20 g/L) with Mycological Peptone (10 g/L) and Micro Agar (12 g/L). *Pseudomonas syringae* pv. *tomato* DC3000 was cultured from a frozen glycerol stock on King Agar B (KB) supplemented with 20 mg/mL peptone protease, 1.5 mg/mL K_2_HPO_4_, 1.5 mL/mL glycerol, 1.5 mg/mL agarose, 25 µg/mL rifampicin, and 5 mM MgSO_4_. The culture was incubated at 28°C in the dark for three days before inoculum preparation.

### 2.9 Hydrogen peroxide quantification

Four mm diameter leaf discs from four-week-old *A. thaliana* plants and five-week-old *S. lycopersicum* ‘Minibel’ were used to determine hydrogen peroxide (H_2_O_2_) production after treatment with flg22 (1 μM) or MIPE (2 μg/mL). Sterile distilled water was used as a mock. H_2_O_2_ production was measured by determining the luminescence produced by the luminol-peroxidase reaction in a Varioskan Lux luminescence reader (Thermo Scientific, Waltham, MA, USA) ^59^. Six biological replicates (considering each leaf disc collected from a single adult plant as one biological replicate) were considered for each experiment. The experiment was performed three times with similar results.

### 2.10 Immunoblot analysis for MAPK activation

*Arabidopsis* seedlings (10-day-old) grown in liquid medium were treated with flg22 (1 μM) or MIPE (2 μg/mL) for 5, 10 and 20 min and frozen with liquid nitrogen. Sterile distilled water was used as mock. Protein extraction and detection of activated MAPKs were performed as previously described ^59^. Briefly, frozen seedlings were homogenized and protein extraction was performed with 50 µL of extraction buffer containing 25 mM Tris-HCl pH 7.8, 75 mM NaCl, 15 mM Egtazic acid (EGTA), 10 mM magnesium chloride, 15 mM sodium β-glycerophosphate pentahydrate, 15 mM bis (4-nitrophenyl) phosphate, 1 mM 1,4-dithiothreitol, 1 mM sodium fluoride, 0.5 mM sodium orthovanadate, 0.5 mM phenylmethylsulfonyl fluoride, 0.1% (v/v) Tween 20, and protease inhibitor cocktail (#P9599; Sigma). Total proteins were quantified using Bradford reagent (Bio-Rad, Hercules, CA, USA). Proteins were separated by SDS-PAGE and transferred onto nitrocellulose membranes which were blocked with Protein-Free Blocking Buffer (TBS; Thermo Scientific) for 2 h at room temperature in agitation. The membranes were incubated overnight in agitation at 4°C with Phospho-p44/42 MAPK (Erk1/2) (Thr202/Tyr204) (4370; Cell Signaling Technology Danvers, MA, USA) diluted 1:1000 in TBS. Then, membranes were washed three times with Tris-Buffered Saline that contains 0.1% (v/v) Tween 20 and then incubated with horseradish-peroxidase-goat anti-rabbit polyclonal secondary antibody (10035943; Thermo Fisher Scientific, Waltham, MS, USA) diluted 1:250 in TBS. Membranes were developed using ECL Western Blotting Substrate (Thermo Scientific, Waltham, MA, USA). Additionally, the membranes were stained with Ponceau S solution to evaluate equal loading. Experiments were conducted three times with similar results.

### 2.11 Gene expression analysis

Ten-days-old *Arabidopsis* seedlings grown in liquid MS/2 medium were treated for 1 h with flg22 (1 μM) or MIPE (2 μg/mL). Sterile distilled water was used as a mock. Seedlings were homogenized in liquid nitrogen. *B. cinerea*-infected or mock-inoculated leaves were frozen in liquid nitrogen. Tissues were homogenized using the mixer mill MM301 (RETSCH GmbH, Haan, Germany) and using stainless steel beads (5 mm in diameter) for about 1 min at 30 Hz. Total RNA was extracted with RNA isolation NucleoZol reagent (Macherey-Nagel, Düren, Germany) according to the manufacturer’s instructions. RNA was treated with RQ1 DNase (Promega, Southampton, UK), and first-strand complementary DNA (cDNA) was synthesized using ImProm-II reverse transcriptase (Promega, Southampton, UK). Quantitative reverse transcription–polymerase chain reaction (qRT-PCR) analyses were performed as previously reported ^60^. Quantitative Reverse Transcription PCR analysis was performed using a CFX96 Real-Time System (Bio-Rad, Hercules, CA, USA). One microliter of cDNA (corresponding to 50 ng of total RNA) was amplified in 20 μL of reaction mix containing 1 × Go Taq qPCR Master Mix (Promega, Southampton, UK) and 0.5 μM of each primer. The conditions for amplification were 95 °C for 2 min; 46 cycles of amplification: 95 °C for 15 s, 58 °C for 15 s, and a final extension of 72 °C for 15 s. Primer sequences were generated with Primer3 software (https://primer3.ut.ee/) (Supplemental Table S1). The expression levels were determined using a modification of the Pfaffl method ^61^ as previously described ^4^. For each sample, the relative expression of each gene was determined by calculating the difference in CT values (ΔCT) between the target gene and the mean of CT value of reference genes *Ubiquitin 5* (*UBQ5*) and *Tubulin 4* (*TUB4*). Three biological replicates (considering each well of the plate with ten seedlings per well as one biological replicate) were performed. The experiment was repeated three times with similar results.

### 2.12 Arabidopsis infection assay with *Botrytis cinerea*

Conidia of *B. cinerea* were harvested by washing the surface of the mycelium with sterile distilled water. Conidia suspensions were filtered in sterile condition to remove residual mycelium, and the conidia concentration was determined using a Thoma cell counting chamber. Four-week-old plants were sprayed with 2 mL of flg22 (1 μM) or MIPE (2 μg/mL) for the protection assay. Distilled water was used as a mock. After 24h, fully developed leaves were infected with 1 × 10^6^ conidia/mL incubated in Potato Dextrose Broth (PDB) at 24 g/L. Six droplets of spore suspension (5 μL each) were placed on the surface of each leaf. Negative control was performed using PDB. Plants were incubated at 24 °C with a 12 h/12 h day/night photoperiod. The priming effect of pretreatment was evaluated by RNA extraction from infected leaves collected at 8 hours post infection (hpi) as described below. The lesion size produced by *B. cinerea* was determined by measuring necrotic tissues using ImageJ software at 48 hpi and was evaluated as an indicator of susceptibility to the fungus. Lesion size is expressed as a mean of a total of 50 lesions and the experiment was repeated three times with similar results.

### 2.13 Arabidopsis and tomato infection assay with *Pseudomonas syringae*

Infection assay of Arabidopsis and tomato with *P. syringae* pv. *tomato* DC3000 was performed as previously reported ^5^. Briefly, four-week-old Arabidopsis and five-week-old tomato plants were sprayed with 2 mL of flg22 (1 μM) or MIPE (2 μg mL^−1^) containing 0.05% Tween 24 MBAL (Croda, Snaith, UK) for Arabidopsis pretreatment, and 2.5% Tween 24 MBAL and 2.5% UEP-100 (Croda, Snaith, UK) for tomato pretreatment as adjuvants. Adjuvant solutions were used as mocks. Bacterial infection was performed 24 h after pretreatments. In Arabidopsis, the priming effect was evaluated by collecting infected leaves at 8 hpi. Next, Arabidopsis and tomato leaf discs were collected from four different plants at 0 (3 hpi) and 3 days post-infection (dpi), and bacteria colonies inside were determined as colony-forming units (cfu) per foliar area. Six biological replicates (considering each serial dilution of bacteria collected from a single adult plant as one biological replicate) were used. The experiment was repeated three times with similar results.

### 2.14 Data analysis

Data were presented as mean ± standard deviation (SD) or standard error (SE) as indicated in the figure legends. The significant differences were evaluated by Student’s t-test or ANOVA analysis followed by Tukey’s test (p ≤ 0.05), as indicated in the figure legends.

## 3. RESULTS

### 3.1 Two-phase olive pomace digestate is enriched in mineral nutrients and characterized by low heavy metal content

To evaluate the potential of two-phase olive pomace digestate for soil enrichment and its suitability as a fertilizer, a comprehensive biochemical characterization was conducted on raw digestate (RD) derived from a biodigester fed exclusively with two-phase olive pomace. The digestate exhibited a pH of 8.0 ± 0.14, a key parameter influencing soil compatibility and plant growth. The organic dry matter content was 78.09 ± 0.02%, while the C/N ratio was 8.0 ± 0.5, indicating a favourable nutrient profile. The dry residue at 105°C, representing the digestate inorganic and non-volatile fraction and providing insights into its stability and impact on soil composition, was measured at 6.69 ± 0.04%. In dry matter, potassium was the most abundant macronutrient at 6.8%, followed by nitrogen at 4.9%, with ammonium contributing 2.0%, and smaller amounts of phosphorus, sulfur, and magnesium (<1%) (Figure 1A). Trace amounts (<0.01% each) of copper, boron, manganese, zinc, and nickel were identified (Figure 1B). Regarding heavy metal content, traces of hexavalent chromium, mercury, lead, and cadmium were found (<0.00015% each) (Supplemental Figure S1A). Unlike other digestates, human pathogenic bacteria, such as *Salmonella* spp., were not detected. These significant findings highlight the potential of two-phase olive pomace digestate as a safe and valuable source of organic matter and nutrients for agricultural applications without significant risk of contamination.

**Figure 1.**
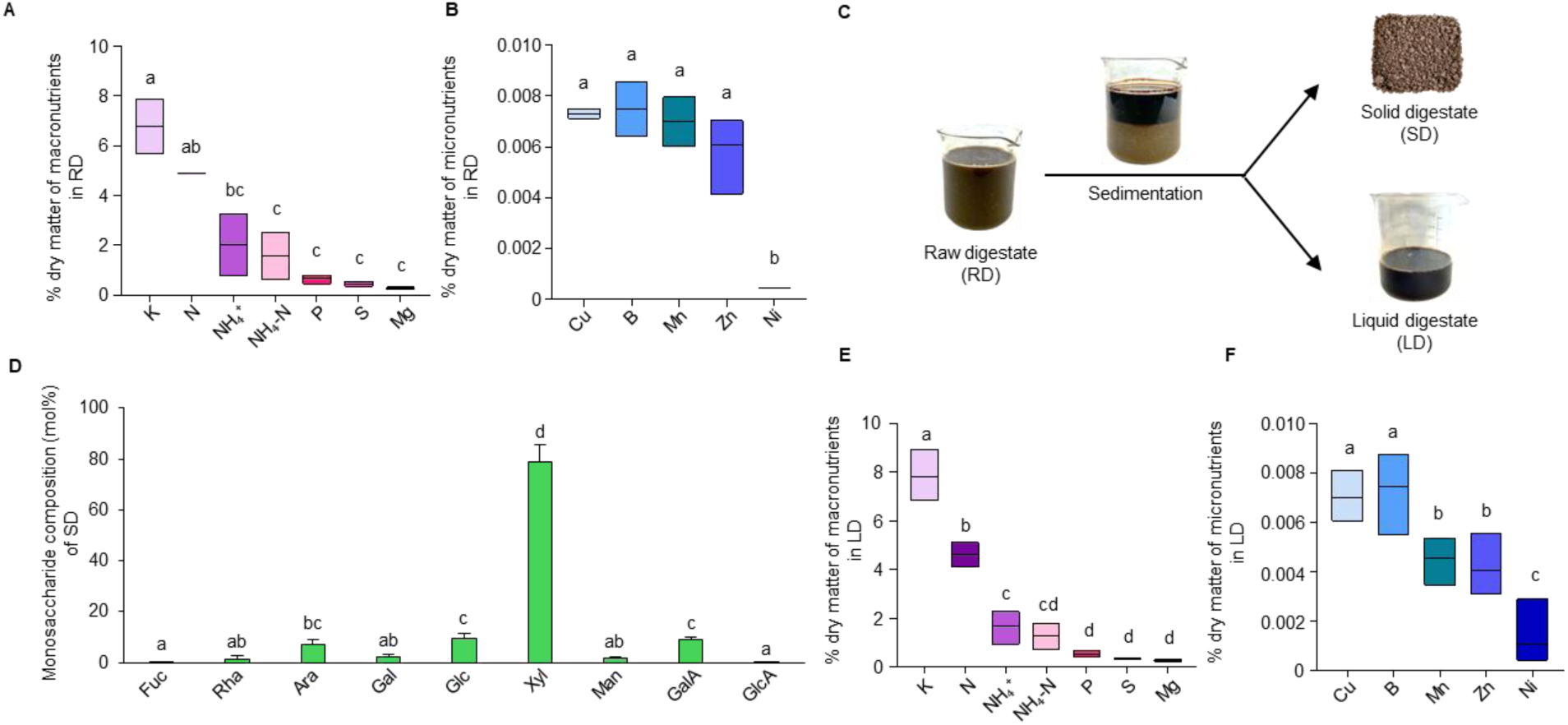
Digestate characterization and representation of chemical composition of the two-phase olive pomace raw digestate and liquid digestate. **(A-B)** Distribution of macro- and micro-nutrients in raw digestate (RD) (% dry matter/raw digestate dry matter). **(C)** Fractionation of olive pomace RD by sedimentation in solid digestate (SD) and liquid digestate (LD). **(D**) Monosaccharide compositions of SD. The molar percentages (mol %) of Fucose (Fuc), Rhamnose (Rha), Arabinose (Ara), Galactose (Gal), Glucose (Glc), Xylose (Xyl), Mannose (Man), Galacturonic Acid (GalA), and Glucuronic Acid (GlcA) were quantified. Results represent the mean ± SD (n=3). (E-F) Distribution of macro- and micro-nutrients in LD. Data represent the mean ± standard deviation (n ≥ 3). The different letters above the boxplots indicate significantly different datasets according to ANOVA followed by Tukey’s test (*p* ≤ 0.05). Potassium (K), Nitrogen (N), Ammonium (NH₄⁺), Ammonium nitrogen (NH₄-N), Phosphorus (P), Sulfur (S), Magnesium (Mg), Copper (Cu), Boron (B), Manganese (Mn), Zinc (Zn), Nickel (Ni).

One of the objectives of this study was to identify which fraction of the digestate may have a greater impact on plant growth. The RD was further characterized by separating it into two primary components: solid digestate (SD) and liquid digestate (LD) (Figure 1C). The SD comprised 32.15 ± 4.3% of the RD (w/w), while the LD accounted for 68.5 ± 4.4% (LD fresh weight/RD fresh weight). Since the cell wall polysaccharides in waste biomasses can influence their effectiveness as soil conditioners, these were extracted (as alcohol insoluble solids) and characterized^55,62^. The sugar component represented 80.2 ± 9.7% of SD dry weight. A detailed chemical characterization of the monosaccharide composition of SD was conducted using HPAEC-PAD (Figure 1D). SD was characterized by a high xylose content (∼80%), followed by low levels (<10% each) of glucose, galacturonic acid, arabinose, mannose, galactose, and rhamnose, along with trace amounts of glucuronic acid and fucose. The high level of xylose indicates that SD is enriched in hemicelluloses (xylans) and contains low levels of pectins. Next, the chemical characterization of LD was performed (Figure 1E-F). The chemical parameters and nutrient composition of LD were similar to those of RD. In particular, the LD exhibited a pH of 8.0 ± 0.08. As shown in Figure 1E, the dry matter of LD contains 7.81% potassium, 4.61% nitrogen, and 1.68% ammonium, the most concentrated components. Phosphorus, sulfur, and magnesium were revealed in smaller quantities (<1%). Trace amounts (<0.01% each) of copper, boron, manganese, zinc, and nickel were also detected. Heavy metals were found in traces (<0.00015%) (Supplemental Figure S1 B). A low percentage of sugars was detected in LD (0.018 ± 0.002% w/v LD). Monosaccharide composition analysis indicated that LD is mainly composed of galacturonic acid (∼30%), rhamnose (∼21%), and glucose (∼18%), with minor amounts of arabinose, galactose, mannose, xylose, fucose, and glucuronic acid (Supplemental Figure S1C).

### 3.2 Soil amendment with olive pomace digestate positively influences plant growth

The effects of RD as a soil amendment and fertilizer were evaluated in a dose-response experiment. Arabidopsis plants were grown in soil amended with RD at 45, 90, or 180 g/kg. After two weeks, rosette fresh weight was measured and compared to plants grown in water-soaked soil used as control (Supplemental Figure S2A). The 45 g/kg dose increased growth by about 25%, while higher doses (90 and 180 g/kg) produced only a 10% increase, yielding a maximum enhancement of 36%. To assess the biostimulant potential under field conditions, the study was extended to *S. lycopersicum* (tomato). A similar pattern was observed in tomato grown in soil amended with RD at 20, 40, or 80 kg/m². After four months, tomato height was measured and compared to the control plants grown in water-soaked soil (Supplemental Figure S2B). The 20 kg/m² dose increased height by approximately 49%, while higher doses led to smaller increases of around 26%, resulting in a total growth increase of 75%. Overall, these findings showed that higher concentrations diminished the growth response, making 45 g/kg and 20 kg/m² optimal for further study on Arabidopsis and tomato, respectively. The next step was to compare the biostimulant potential of RD with solid digestate (SD) and liquid digestate (LD). Arabidopsis plants were grown in soil amended with RD (45 g/kg), SD (15 g/kg), or LD (30 g/kg). After two weeks, rosette fresh weight was measured and compared to a control. All treatments exhibited a significant stimulatory effect on Arabidopsis growth (Figure 2A). Specifically, a 43% increase in growth was observed in Arabidopsis plants cultivated in soil amended with RD compared to the unamended control. Among the tested fractions, SD induced a significant 57% growth increase, while LD exhibited a growth enhancement of only 12% compared to the control. The study was extended to tomato, cultivating plants in soil amended with RD (20 kg/m²), LD (14 kg/m²), or SD (6 kg/m²) fractions. The growth parameters were quantified after four months and compared to untreated controls (Figure 2 B-C). RD amendment significantly increased height and fruit number compared to untreated plants (34% and 232%, respectively). Both digestate fractions enhanced tomato height and productivity compared to the control. As observed in Arabidopsis, a greater increase in plant height and fruit number was noted for plants grown in soil amended with SD (approximately 63% and 315%, respectively) compared to those amended with LD (19% and 150%, respectively). These results indicate that two-phase olive pomace digestate can act as a biostimulant for Arabidopsis and tomato growth and fruit production, with SD proving more effective in stimulating plant growth compared to LD.

**Figure 2.**
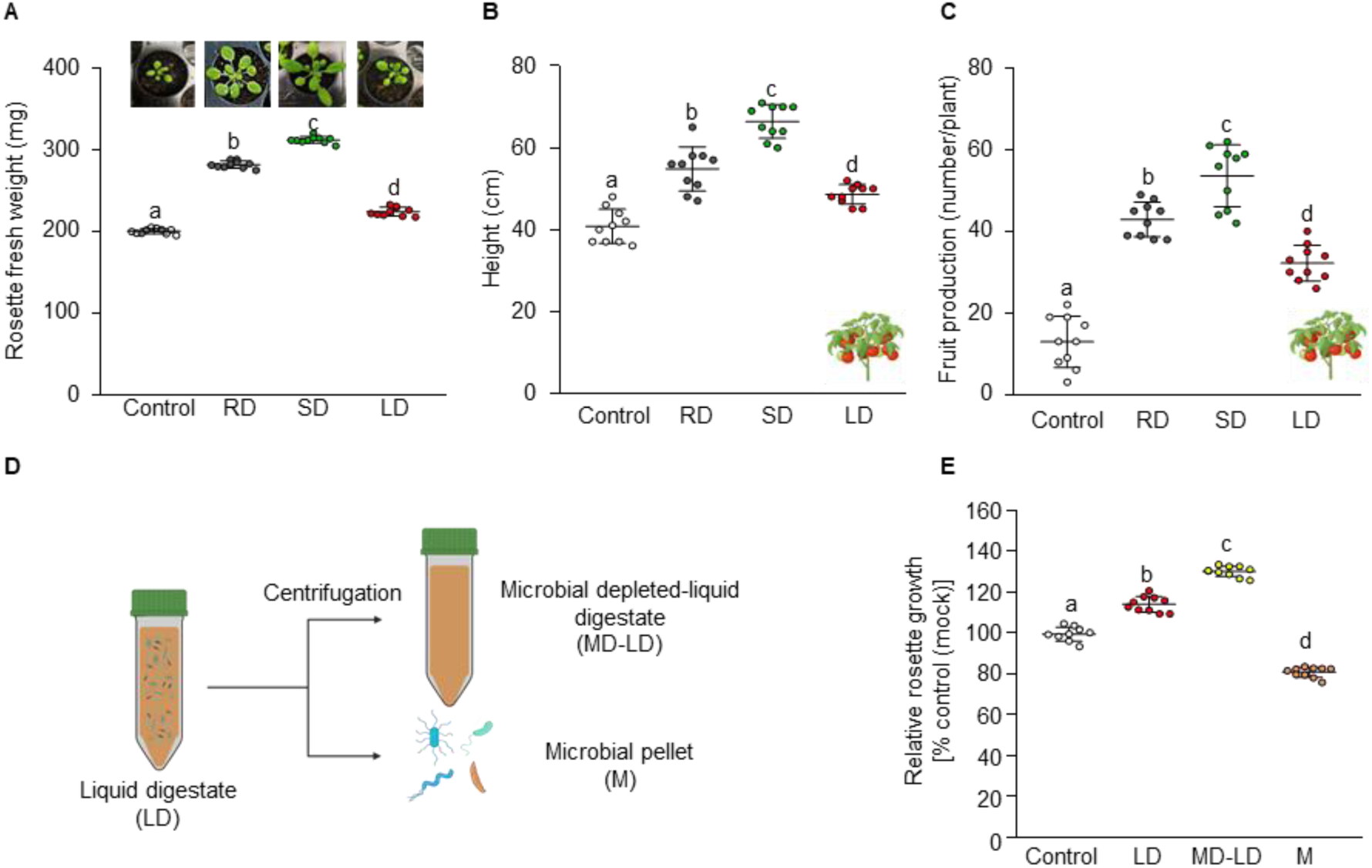
Different fractions of olive pomace digestate improved plant growth. (**A**) Effects of soil amended with RD (45 g/kg), SD (15 g/kg) or LD (30 g/kg) fractions on *A. thaliana* growth. (**B**-**C**) Effects of soil amended with RD (20 kg/m^2^), SD (6 kg/m^2^) or LD (14 kg/m^2^) on height (**B**) and fruit production (**C**) of *S. lycopersicum,* grown on field. (**D**) Reduction of microbial load in liquid digestate through centrifugation, resulting in a microbe-depleted liquid digestate (MD-LD) solution. (**E**) Effects of soil amended with LD, MD-LD (both 30 g/kg), or M (1.5 g/kg) on *Arabidopsis*. The control (mock) represents plants grown in water-soaked soil. The value are expressed as percentage relative to mock. Data shown represent the mean ±SD (n=10). The experiments were repeated three times with similar results. The different letters on the bars indicate significantly different datasets according to ANOVA followed by Tukey’s test (p ≤ 0.05).

### 3.3 Microbial fraction reduces digestate biostimulant efficiency

We hypothesized that specific components in the LD fraction might reduce its bio-stimulatory potential. The presence of galacturonic acid-based carbohydrates in LD (Figure 1D) suggests a possible trade-off between growth and defense responses induced by OG. To explore this hypothesis, pectic fragments were precipitated from LD using ethanol fractionation, and the oligosaccharide profile was analysed using HPAEC-PAD ^5,57^. However, no OG or other carbohydrate-based elicitors peaks were detected in LD (Supplemental Figure S3A-B).

Given that many protein-derived MAMPs are shared across various microbial species and considering that digestate can be enriched with diverse bacterial and fungal populations as well as residual plant biomass, we hypothesized that digestate may function as a continuously proliferating, low-cost reservoir for MAMPs/DAMPs-based phytovaccines. To test this assumption, LD was centrifuged generating two fractions: a microbial-depleted LD (MD-LD) and a microbial-enriched pellet (M) (Figure 2D). Arabidopsis plants were then cultivated in soil amended with varying concentrations of LD, MD-LD (both at 30 g/kg), or M (1.5 g/kg), with the amount of M corresponding to the relative proportions found in the original LD. After two weeks, the rosette fresh weight was measured and compared to the control (Figure 2E). Notably, an increase in Arabidopsis growth was observed in plants grown in both LD- and MD-LD-amended soil. The growth increase was significantly higher in the MD-LD treatment compared to the LD, with a difference of approximately 16.7%. Conversely, the M treatment led to a decrease in plant growth by about 19.2 %. These data indicate that the microbes present in LD negatively affect its fertilizing potential on both Arabidopsis and tomato growth.

### 3.4 Several bacterial and fungal species were identified in the liquid fraction of olive mill digestate

Detailed information on the microbial component isolated in the M fraction was obtained through taxonomic characterization of the microbial community using 16S and ITS rRNA metabarcoding to identify bacteria and fungi populations, respectively. This analysis revealed a complex bacterial genus-level community structure, with nearly 50% of sequenced reads assigned to uncultured organisms (Figure 3A). Other predominant genera (relative abundance >3%) included *Luteimonas*, *Planomicrobium*, *Caldicoprobacter*, *Pseudomonas*, and *HN-HF0106*. Sequence data further confirmed the presence of anaerobic bacteria such as *Tissierella* and *Sedimentibacter* (*Peptostreptococcaceae*) and genera within the *Ruminococcaceae* family (*UCG-010* and *Ruminiclostridium*). Notably, most uncultured organisms were associated with the *DTU014* (nearly 40%), followed by *Chloroflexi* (22%), *Firmicutes* (14%), *Collierbacteria* (6%), *SAR324*, and *Bacillaceae* (3%) (Figure 3B). Taxonomic assignment of the fungal community was challenging, likely due to the low DNA concentration in the samples. This limitation may explain why 71% of the sequences were identified as *Fungi gen. incertae sedis*, indicating undefined broader taxonomic relationships. Despite the low abundance of ITS sequences, notable findings included 7% of the sequences assigned to the genus *Pichia* and *Basidiomycota,* and >3% assigned to *Paraglomerales*, *Brettanomyces*, and *Glomus*. These findings indicate that the microbial community in the digestate is highly diverse, including both bacteria and fungi populations.

**Figure 3.**
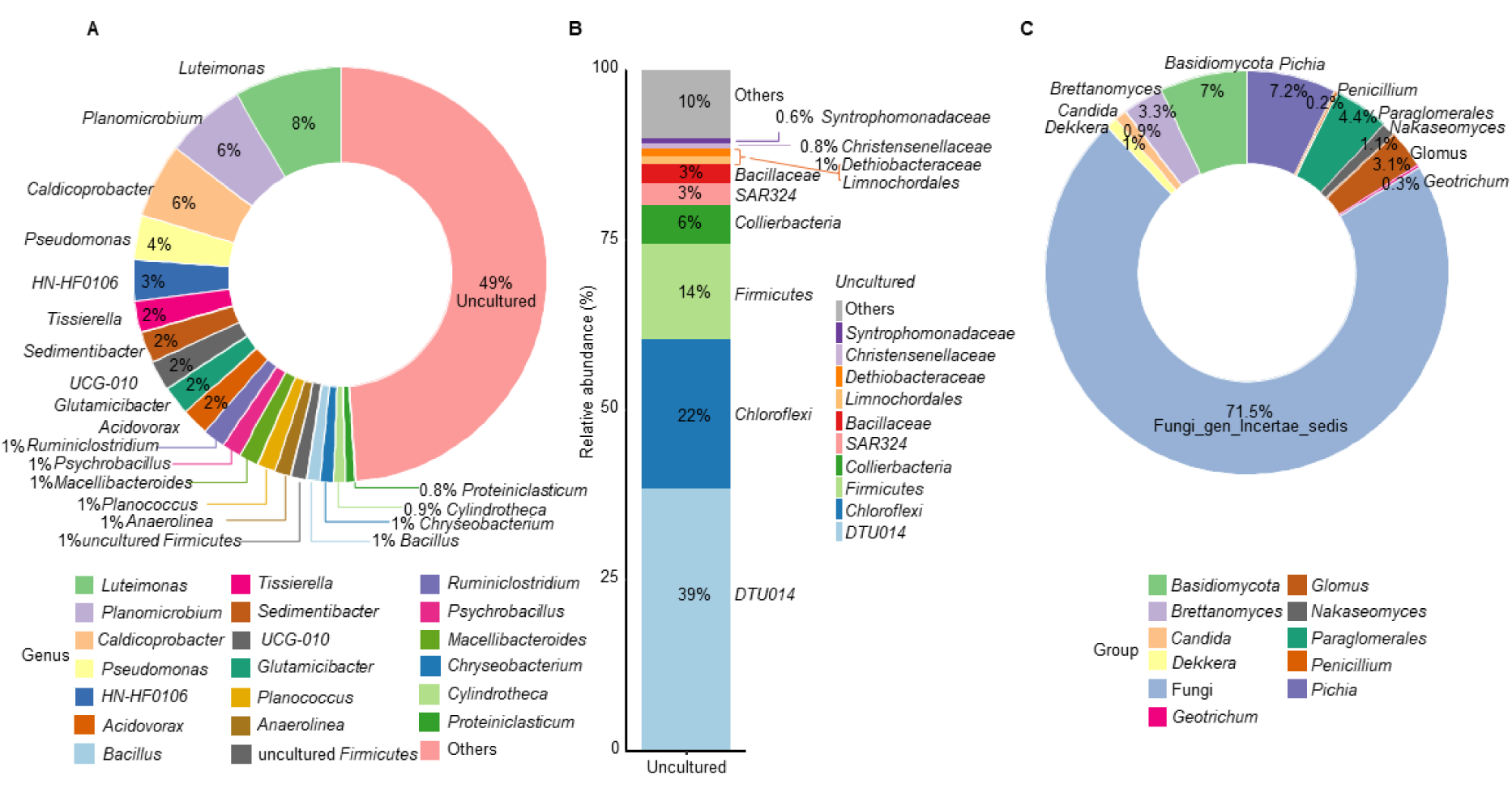
Abundance analysis of 16S rDNA of bacterial cells and rDNA internal transcribed spacer (ITS) region of fungal cells from digestate. (**A**) Donut chart describing the proportion of reads assigned to each genus of bacterial community. (**B**) Bar plot showing the composition of the «uncultured» fraction of bacterial community. Each percentage is calculated considering a restricted dataset, represented by the hits classified as «uncultured» at genus level (**C**) Donut chart describing the proportion of reads assigned to each genus of fungal community, represented with different colours. Percentage of abundance is indicated by the integer within each slice.

### 3.5 Immunogenic proteins are present in olive pomace digestate fraction

Next, we hypothesized that digestate can serve as a cost-effective source of MAMPs/DAMPs. The extraction of proteinaceous elicitors from a LD was attempted. M fraction was sonicated, and the protein extract obtained was referred to as Microbial-enriched Protein Extract (MIPE). The yield of MIPE extract was 20 mg/L of digestate liquid fraction. The presence of nucleic acids was excluded by spectrophotometric analysis (about 10 ng/μL) and gel electrophoresis (Supplemental Figure S3C). Next, a proteomic approach using mass spectrometry was employed to identify the proteins in MIPE. For protein identification, raw proteomic data were queried against the Uniprot database (https://www.uniprot.org/), which includes proteins from bacteria, fungi, and the olive plant identified during taxonomic characterization. A total of 1,172 proteins were identified. The dataset was then filtered to include only those proteins consistently detected in at least two out of three biological replicates, yielding a refined set of 67 robustly identified proteins. The proteomic analysis indicated that most of the proteins belong to the bacterial community (50.7%), followed by fungi (40.3%), and olive plant (8.9%) (Figure 4 A). Functional Gene Ontology (GO) assignation was undertaken to provide an overview of the functional composition of protein identified in MIPE. The identified proteins were grouped into 7 broad functional classes, which were manually compiled by grouping GO terms related to different biological processes. These GO terms were sourced from Gene Ontology database. The entire dataset of proteins was first categorized by biological process. Then, a detailed analysis was carried out by considering individually the three components accountable for the reserve of proteins: bacteria, fungi and olive (Figure 4 B). In bacteria, the identified proteins were mainly associated with the biological process involved in interaction with the host (35.7%), followed by cellular component organization or biogenesis (29.2%), metabolic process (16.7%), cellular localization (8.3%), response to stimulus, and cell cycle process (both 4.2%). In fungi, metabolic process was the most representative (64.3%), followed by biological process involved in interaction with the host (21.4%), cellular component organization or biogenesis, and cellular localization (both 7.1%). Among the proteins with olive origin, there were proteins related to defense response, metabolic processes, and cell cycle processes (all three 33.3%). The association of proteins identified in MIPE with categories of biological processes involved in interaction with host and defense response suggested the presence of proteins that could have a pivotal role in plant-pathogen interaction. The analysis indicates that MIPE contains proteins with considerable size variation (Figure 4C). A significant proportion consists of proteins with a molecular weight corresponding to <300–400 amino acids, suggesting the presence of small peptides with potential elicitor activity. Additionally, proteins with high molecular weight were also identified.

**Figure 4.**
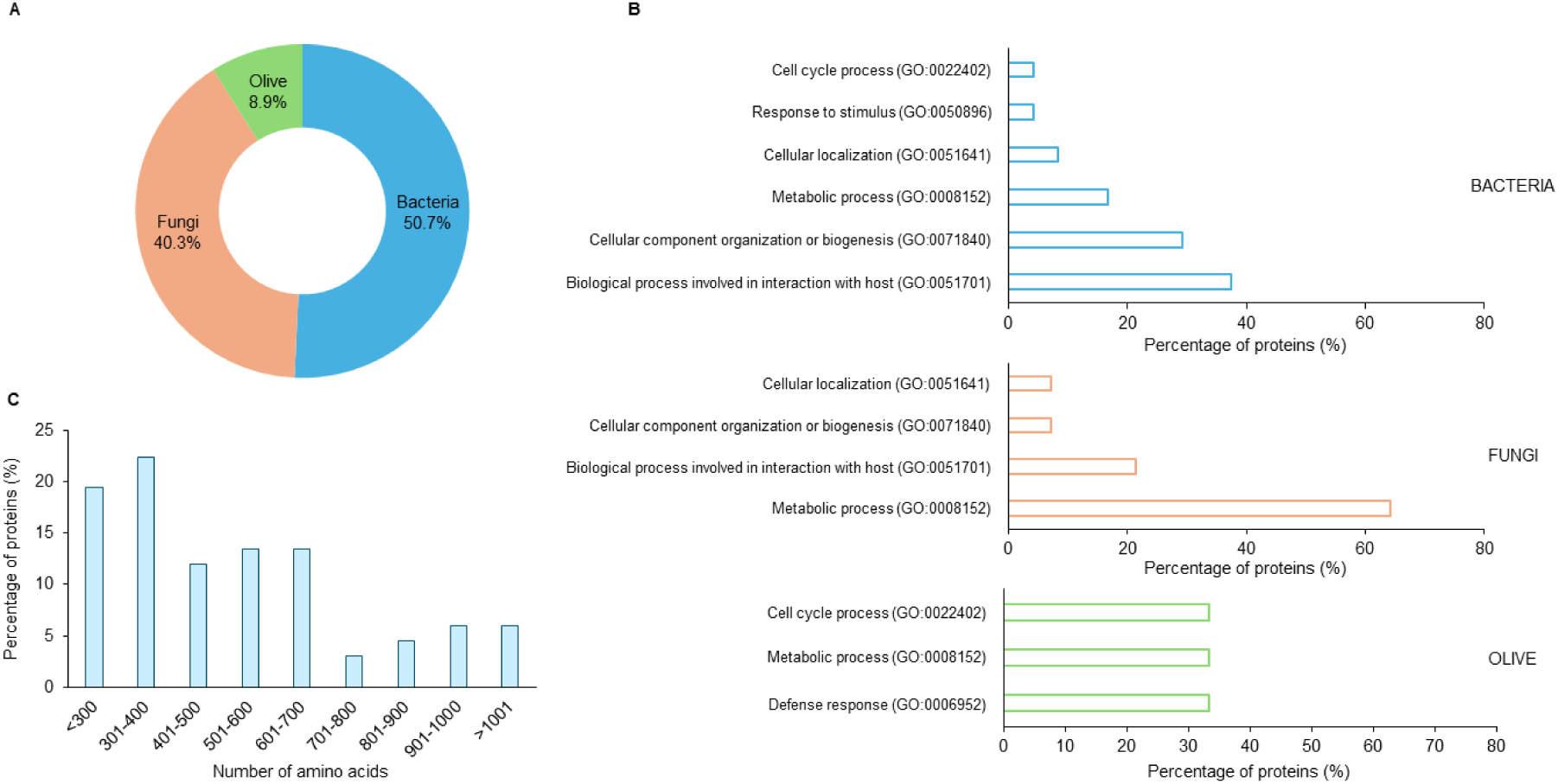
Protein analysis in MIPE extract. (**A**) Donut chart describing the proportion of protein assigned to bacteria, fungi or olive, represented with different colours. Percentage of abundance is indicated by the integer within each slice. (**B**) GO term category of biological processes of identified proteins sorted by bacteria, fungi, and olive groups. (**C**) Amino acids length of identified proteins.

Intriguingly, microbial functional proteins involved in plant immune responses were detected (Table 1). In particular, bacterial MAMPs were observed, including elongation factor Tu, releasing a fragment of 18 amino acids (elf18) that can be recognized by Elongation Factor Tu Receptor (EFR) in Arabidopsis ^63^ and flagellin, whose immunogenic peptides can be perceived via Arabidopsis Flagellin Sensitive 2 (FLS2) ^64^. Enzymatic MAMPs derived from bacterial and fungal communities were also detected, including endo-1,4-beta-xylanases A, pectate lyases, and histidine kinases whose elicitor activities can be independent of their enzyme activities ^65–67^. A homolog to golven 1-2A, a phytocytokine known as inducible peptidic DAMP, was also identified ^38,68^. The proteomic analysis also showed enzymes that could produce signaling molecules from hemicellulose components of the plant cell wall, such as xylanases, and from plant cell wall pectin like rhamnosidases and pectate lyases. ^69–73^.

**Table 1.** List of proteins and peptides identified in MIPE, known to act as MAMPs, phytocytokines, or enzymes that release DAMPS in plant immunity.

| MAMPs | Origin | Reference |
| --- | --- | --- |
| Elongation factor Tu | Bacteria | 31,63 |
| Flagellin | Bacteria | 64,114,127,128 |
| Endo-1,4-beta-xylanase A | Bacteria/Fungi | 65,129,130 |
| Pectate lyase | Bacteria/Fungi | 67,131 |
| Histidine kinase | Bacteria/Fungi | 66 |
| Phyto cytokines |  |  |
| Homolog to GOLVEN 1-2 | Olive | 38,68 |
| Enzymes potentially releasing DAMPs |  |  |
| α-L-rhamnosidase | Fungi | 70 |
| Endo-1,4-β-xylanase A | Bacteria/Fungi | 69,72,73 |
| Pectate lyase | Bacteria/Fungi | 118,132,133 |

### 3.6 MIPE induced PTI hallmarks in Arabidopsis plants

Next, we examined whether MIPE can act as an elicitor of plant immune responses. To determine a concentration that would enhance immunostimulation without any defence-growth trade-off ^43,74^, a dose-response analysis of MIPE treatment was firstly performed on Arabidopsis seedling growth. Arabidopsis seedlings were cultivated in a liquid medium supplemented with MIPE at different concentrations (1, 10, or 100 μg/mL) and the seedling weights were quantified after 5 days of treatment (Supplemental Figure S2C). Seedlings treated with MIPE at 1 and 10 μg/mL showed a growth not significantly different from mock-treated seedlings. Instead, the addition of MIPE at 100 μg/mL produced about 35% of seedling growth inhibition. Next, we explored the possibility of MIPE to activate PTI hallmarks in Arabidopsis plants, including hydrogen peroxide (H_2_O_2_) production, activation of MAPK phosphorylation cascades, and upregulation of defense-related genes ^75,76^. The production of H_2_O_2_ was quantified in leaf discs of Arabidopsis plants, after treatment with 1 μg/mL MIPE or water (mock), by luminol-based assays. MIPE induced a transient burst of H_2_O_2_ compared to mock treatment (Figure 5 A). The elicitor capacity of MIPE was compared with that induced by 1 μM flg22, used as comparison. Flg22 induced a transient burst of H_2_O_2_ that reached its maximum level 10 minutes after treatment (Figure 5 B). This response was faster compared to the response induced by MIPE, where the peak occurred 20 minutes after treatment. To confirm that MIPE triggers additional PTI responses, we performed an immunoblot analysis to evaluate the temporal phosphorylation patterns of MAPKs (MAPK3, MAPK6, MAPK4/11) in mock- and elicitor-treated (1 μg/mL MIPE or 1 μM flg22) Arabidopsis seedlings. MIPE-treated seedlings showed the phosphorylation of MAPK6 and MAPK3 after 5 min and slight MAPK4/11 phosphorylation after 10 min that increased at 20 min (Figure 5 C). Interestingly, the phosphorylation pattern of MIPE was similar to the flg22 one, used as comparison, where however higher phosphorylation of all MAPKs was observed already at 10 min after treatment. Next, we explored the expression of three PTI-reporter genes, *CYP81F2*, *FRK1*, and *WRKY53*, after 1 h of Arabidopsis seedling treatment with MIPE (1 µg/mL), flg22 (1 μM) or mock. *CYP81F2* (*CYTOCHROME P450, FAMILY 81*) encodes a cytochrome P450 monooxygenase involved in the biosynthesis of indole glucosinolates ^77^; *FRK1* (*FLG22-INDUCED RECEPTOR-LIKE KINASE 1*) encodes a leucine-rich repeat receptor kinase that functions in defense signaling pathways ^78^; *WRKY53* (*WRKY DNA-BINDING PROTEIN 33*) encodes a transcription factor contributing to basal resistance against *P. syringae* ^79^. The MIPE or flg22 treatments strongly stimulated the expression of all PTI genes (Figure 5 D-E-F). However, interesting differences arose when comparing the gene induction patterns of MIPE with flg22, used as comparison. Flg22 was more effective in stimulating CYP81F2 expression, while MIPE induced higher FRK1 expression. Both MIPE and flg22 treatments showed similar levels of *WRKY53* expression. Overall, our findings suggest that the exogenous application of specific MIPE concentration to Arabidopsis can induce multiple early immune responses without impacting plant growth.

**Figure 5.**
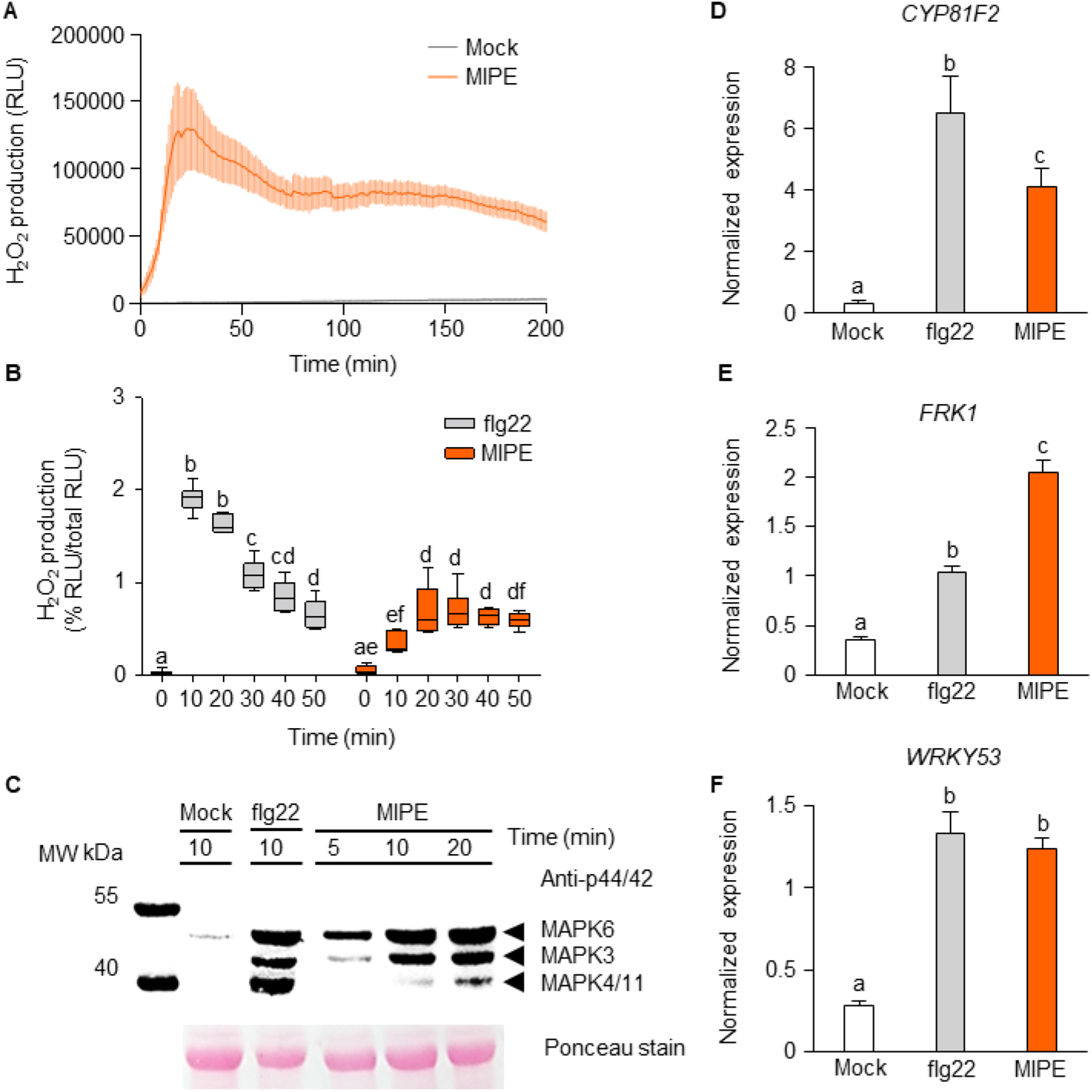
MIPE activated immunity hallmarks in Arabidopsis tissues. (**A**) Kinetics of H_2_O_2_ production measured by Luminol reaction after treatment with sterile distilled water (mock), or MIPE (1 μg/mL) in four-week-old Arabidopsis leaf-discs and reported as Relative Luminescence Units (RLU). Data represent mean ± SE (n=6). The experiments were repeated three times with similar results. (**B**) Ratio of RLU values after flg22 (1 μM), or MIPE (1 μg/mL) at 0, 10, 20, 30, 40, and 50 minutes respect to the total RLU. Data are presented as box plots, with the centre line showing the median, the box limits showing the 25^th^ and 75^th^ percentiles, and the whiskers showing the full range of data. The experiments were repeated three times with similar results. (**C**) MAPK activation in Arabidopsis seedlings in response to mock, flg22 (1 μM), or MIPE (1 μg/mL) treatments. MAPK phosphorylation was determined by western blot using the phospho-p44/42 MAPK antibody at different time points (5, 10, and 20 minutes). Equal loading was confirmed by Ponceau staining. Experiments were conducted three times with similar results. MW =Molecular Weight marker. Expression of *CYP81F2* (**D**), *FRK1* (**E**), and *WRKY53* (**F**) after mock, flg22 (1 μM), or MIPE (1 μg/mL) treatments. The expression of defense genes was analyzed by quantitative RT-PCR at 1h after treatments on 10-days-old Arabidopsis Col-0 seedlings. The expression levels were normalized to *UBQ5* and *TUB4* expression levels. Data represent the mean ±SE (n=3) and the experiment was performed three times with similar results. Different letters indicate significant differences according to ANOVA followed by Tukey’s test (*p ≤* 0.05).

### 3.7 MIPE reduced Arabidopsis and tomato disease symptoms caused by bacterial and fungal pathogens

Following evidence that MIPE can trigger PTI responses, we assessed its potential protective effect as a pretreatment for Arabidopsis against the necrotrophic fungus *B. cinerea*. Four-week-old adult plants were foliar-sprayed with MIPE (1 µg/mL), flg22 (1 μM), or mock, and 24 h later, leaves were inoculated with *B. cinerea* spores (Figure 6A). Primed expression of the PTI genes *CYP81F2* and *PAD3* was evaluated at 8 hours post-infection (hpi) and compared with mock pretreatment (Figure 6 B-C). *PAD3* (*PHYTOALEXIN DEFICIENT 3*) encodes a key biosynthetic enzyme involved in the biosynthesis of the antimicrobial compound camalexin and is essential for elicitor-induced resistance to *B. cinerea* ^80,81^. MIPE- and flg22-pretreated plants showed significant overexpression of *CYP81F2* and *PAD3* genes compared to mock-pretreated plants. Interestingly, induction levels of both genes were comparable between the two elicitors. After 48 hpi, MIPE and flg22 pretreatments significantly improved the resistance of Arabidopsis to *B. cinerea* (Figure 6 D-E). In particular, plants pretreated with MIPE showed a greater reduction of lesion area (about 69%) than flg22 pretreated ones (about 54%) compared to mock.

**Figure 6.**
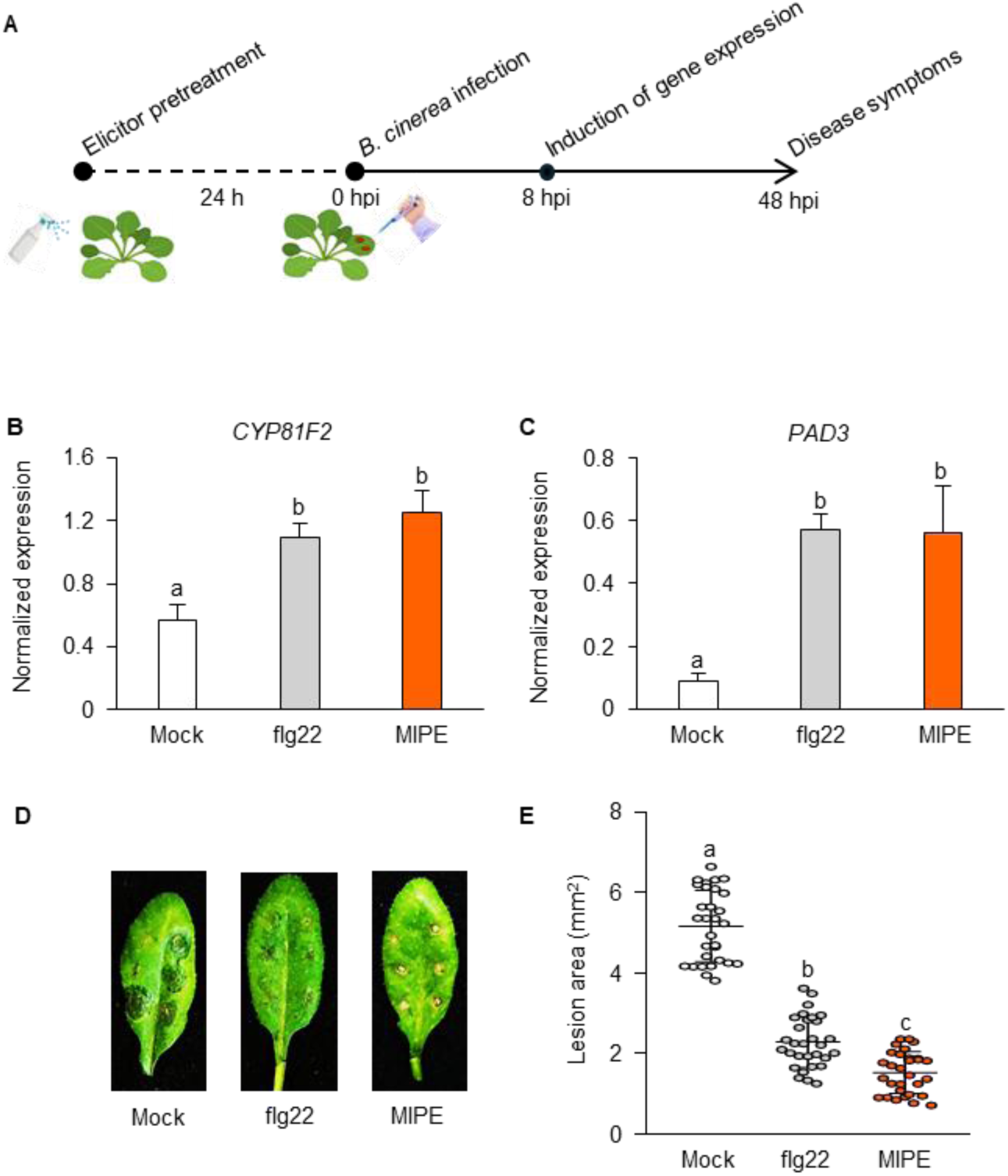
Pretreatment of Arabidopsis with MIPE enhanced immune response and protection against *B. cinerea*. (**A**) Four-week-old Arabidopsis Col-0 leaves were pretreated with mock, flg22 (1 μM), or MIPE (1 μg/mL) and 24 h later were inoculated with *B. cinerea*. (**B**-**C**) Quantitative RT-PCR analysis in *B. cinerea*-infected leaves collected at 8 h post infection (hpi). mRNA expression levels are normalized to *UBQ5* and *TUB4* expression levels. Data represent the mean ±SE (n=3). (**D**-**E**) Lesion areas of *B. cinerea*-infected leaves were measured at 48 h after the inoculation. Values are means ± SD of at least 50 lesions. The experiments were performed three times with similar results. Different letters indicate significant differences according to ANOVA followed by Tukey’s test (*p ≤* 0.05).

The MIPE induction of *WRKY53* expression previously observed in Arabidopsis seedlings suggested a potential priming and protective effect of MIPE against *P. syringae* ^82^. To confirm this hypothesis, adult Arabidopsis plants were pretreated with MIPE (1 μg/mL), flg22 (1 μM), or mock, and after 24 h, they were sprayed with a suspension of *P. syringae* (Figure 7 A). The primed state was analysed by monitoring the expressions of the PTI genes *CYP81F2* and *WRKY53* at 8 hpi and compared with mock pretreatment (Figure 7 B-C). Pretreatment of Arabidopsis with MIPE and Flg22 resulted in a comparable and significant upregulation of WRKY53 and CYP81F2 relative to mock pretreatment. To investigate the ability of MIPE to protect Arabidopsis against *P. syringae*, bacterial quantification was performed at 0 and 3 days post-infection (0 dpi and 3 dpi, respectively) and expressed as the logarithm of colony forming unit (CFU) per leaf area (cm^2^) (Figure 7 D). At 0 dpi, the same number of bacteria was observed in both elicitor-pretreated and mock-pretreated plants, proving that the differential bacterial growth depends solely on the pretreatment. Interestingly, no bacterial growth increase was revealed in the MIPE- and flg22-pretreated plants after 3 dpi, compared with 0 dpi. On the other hand, in mock-pretreated plants, bacterial growth was significantly increased by 39 % after 3 dpi, compared with 0 dpi. We then investigated whether the knowledge acquired from the model species Arabidopsis could be applied to crop species such as tomato. Leaf discs collected from five-week-old tomato plants were treated with MIPE (1 µg/mL), or mock and H_2_O_2_ production was quantified by luminol-based assays (Figure 8 A). MIPE induced a transient increase in H_2_O_2_ levels compared to the mock treatment. H_2_O_2_ production of MIPE treatment was compared with one of flg22 treatment (Figure 8 B). As observed in Arabidopsis, flg22 treatment elicited a transient burst of H_2_O_2_, reaching its peak concentration 10 minutes post-treatment. This response occurred more rapidly than that triggered by MIPE, where the maximum H_2_O_2_ level was observed about 30 minutes after treatment. These results prompted us to investigate possible tomato protection against pathogens. Five-week-old tomato plants were pretreated with MIPE (1 µg/mL), flg22 (1 μM) or mock 24 h before spray-inoculation with *P. syringae* (Figure 8 C). In mock-pretreated plants, bacterial colonies increased by about 21 % after 3 dpi, compared with 0 dpi. As observed in Arabidopsis, flg22 and MIPE pretreatments inhibited bacterial growth as bacterial colonies were very similar to those quantified at T0 dpi (Figure 8 D). Overall, the results indicate MIPE as a new elicitor mixture able to enhance Arabidopsis and tomato immune performance and protect against the pathogens *B. cinerea* and *P. syringae*.

**Figure 7.**
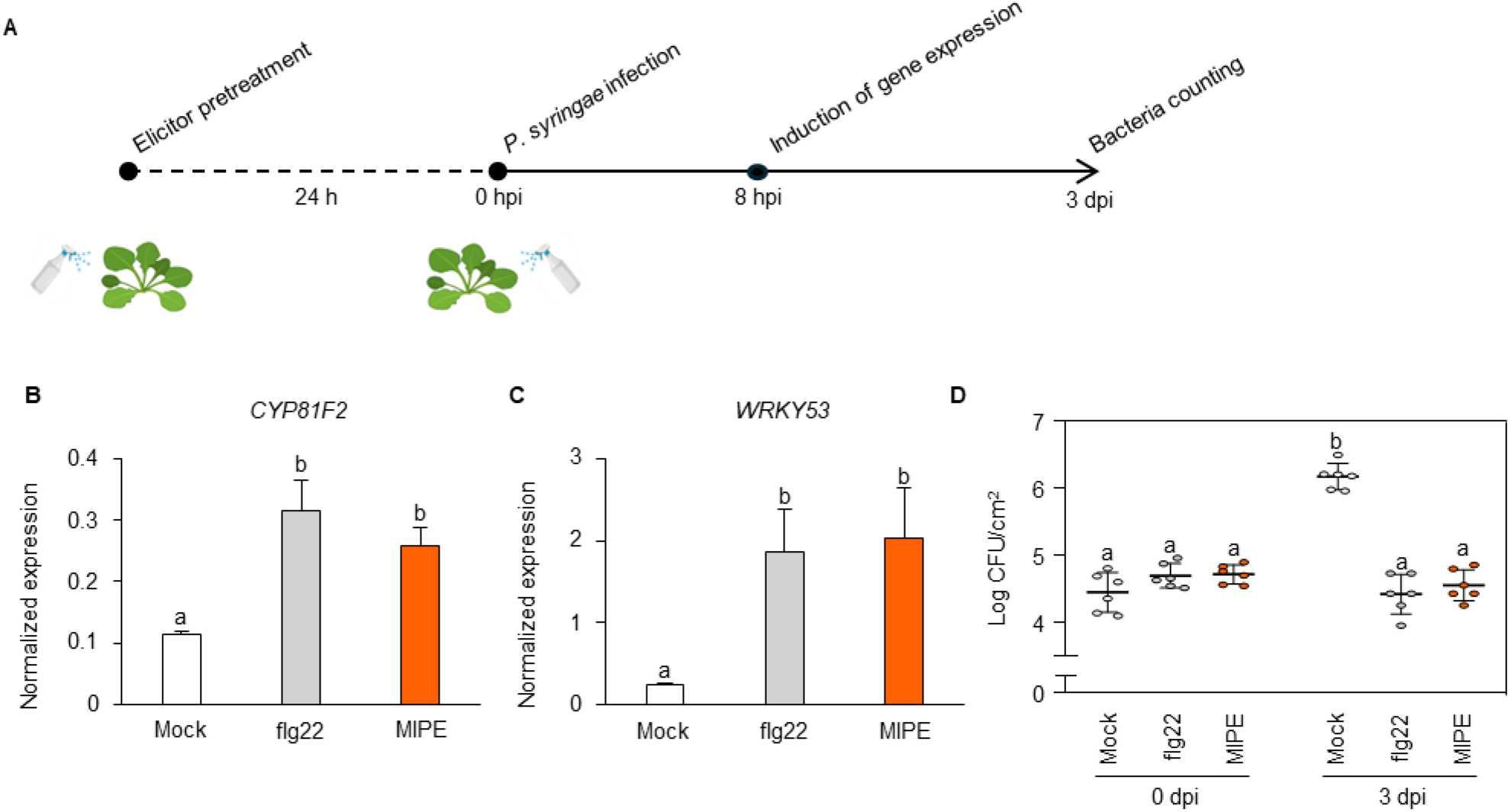
Pre-treatment of Arabidopsis with MIPE enhanced immune response and protection against *P. syringae*. (**A**) Leaves of 4-weeks-old Arabidopsis Col-0 plants were pre-treated with adjuvant solutions (mock), flg22 (1 μM), or MIPE (1 μg/mL) 1 day before bacterial inoculation. (**B**-**C**) Quantitative RT-PCR analysis in *P. syringae*-infected leaves collected at 8 hpi. mRNA expression levels are normalized to *UBQ5* and *TUB4* expression levels. Data represent the mean ±SE (n=3). (**D**) Colony forming units (Log CFU) of *P. syringae* per leaf area (cm^2^) were determined 0 and 3 days post-infection (0 dpi and 3 dpi). Data represent mean ±SD (n=6). The experiments were performed three times with similar results. Different letters indicate significant differences according to ANOVA followed by Tukey’s test (*p ≤* 0.05).

**Figure 8.**
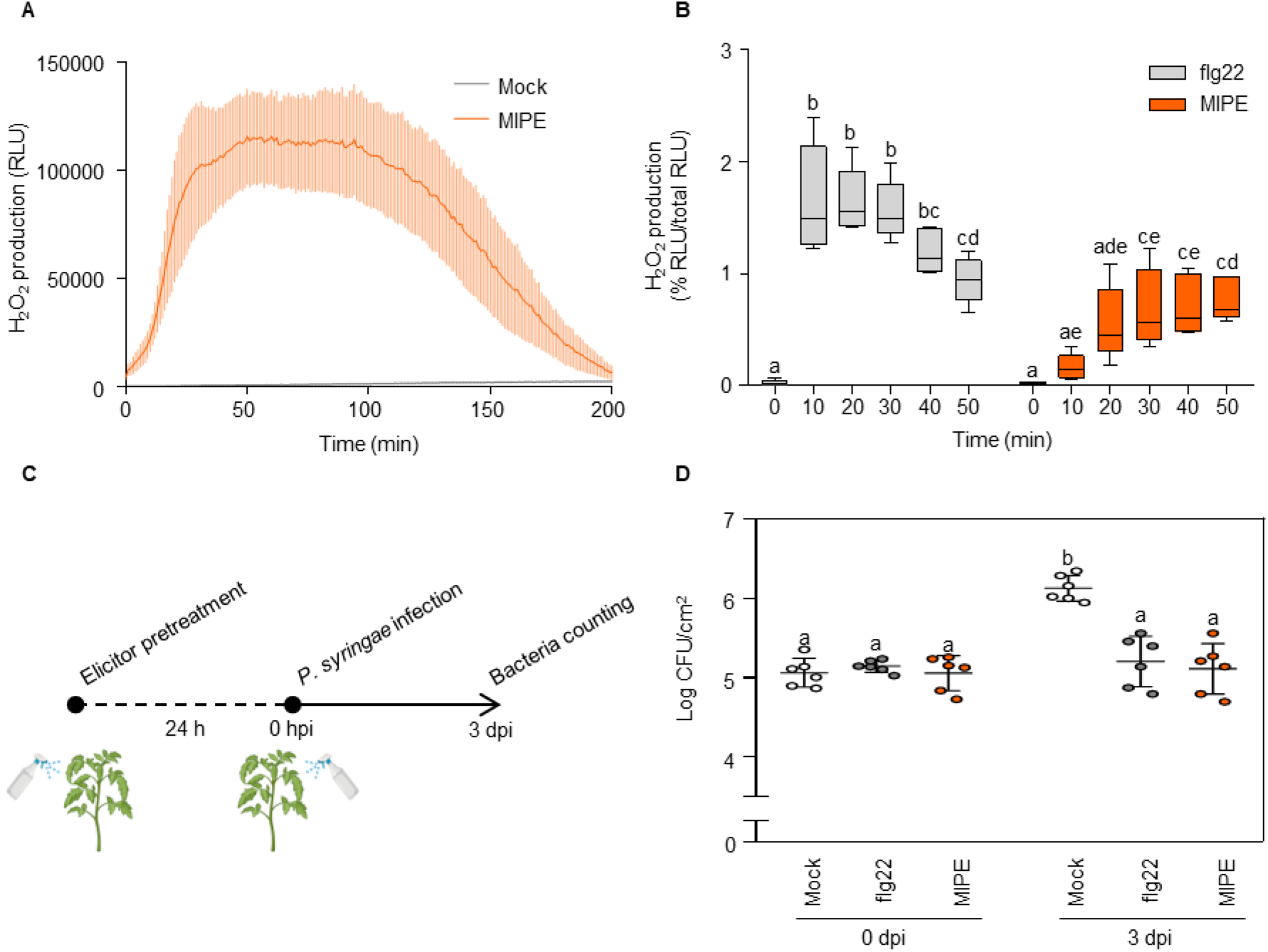
MIPE activated ROS production and induced *P. syringae* pv. *tomato* DC3000 resistance in tomato. (**A**) ROS production by Luminol reaction after treatment with distilled water (mock) or MIPE (1 μg/mL) in five-week-old tomato leaf-discs. Data represent mean ± SE (n=6). (**B**) Ratio of RLU values after flg22 (1 μM), or MIPE (1 μg/mL) at 0, 10, 20, 30, 40, and 50 minutes respect to the total RLU. (**C**) Leaves of five-week-old tomato Minibel plants were pre-treated with adjuvant solutions (mock), flg22 (1 μM), or MIPE (1 μg/mL) 1 day before *P. syringae* inoculation. (**D**) Bacterial CFU per leaf area (cm^2^) were determined at 0 dpi and 3 dpi. Data represent mean ±SD (n=6). The experiments were performed three times with similar results. Different letters indicate significant differences according to ANOVA followed by Tukey’s test (*p ≤* 0.05).

## 4. DISCUSSION

The increasing adoption of biogas plants by olive mills represents a strategic shift toward sustainability, converting olive pomace into bioenergy to reduce emissions and enhance profitability in the olive oil sector. However, the potential of olive pomace digestate’ valorization remains underexplored. This study demonstrates the strong promise of two-phase olive pomace digestate as a biostimulant for Arabidopsis and tomato growth and productivity. Several characteristics make this digestate particularly useful as a soil amendment and fertilizer. Its slightly alkaline pH makes it compatible with various soils and useful for counteracting soil acidification ^83^. The high organic dry matter content emphasizes its richness in organic carbon, vital for improving soil structure, water retention, and supporting microbial activity, positioning it as an excellent soil amendment ^84,85^. The favourable C/N ratio ensures balanced carbon and nitrogen availability, promoting nutrient cycling and making nitrogen readily accessible to plants ^15,86^. The high potassium and ammonium content in two-phase olive pomace digestate enhances its value as a nutrient-rich fertilizer, promoting plant growth and improving soil fertility ^18^. Trace micronutrients such as copper, boron, manganese, zinc, and nickel provide added nutritional value for plants. Two-phase olive pomace digestate used in this study presented low ash content minimizing the risk of soil salinization and toxicity, and the absence of heavy metals and harmful bacteria like *Salmonella* spp. These characteristics offer enhanced safety for agricultural use compared to digestates from the anaerobic digestion of livestock waste, crop residues, and urban waste, which may contain pollutants and pathogens, requiring careful management and sterilization to reduce potential risks ^87–91^.

Nevertheless, several aspects of the agronomic use of olive pomace digestate remain insufficiently investigated, with key factors influencing its efficiency still hidden or poorly understood. This study emphasizes the importance of digestate refinement in enhancing its agricultural performance, shedding light on the unique roles and contributions of its fractions to soil health and crop productivity. Specifically, the two-phase olive pomace RD was separated into SD and LD fractions. Among these, the SD fraction exhibited a stronger impact on enhancing plant growth and productivity compared to the LD fraction. The lower effect exerted by LD does not appear to be related to the mineral nutrient content, as it is similar to that observed in RD. We also ruled out the possibility that the reduced fertilizer efficacy of LD is attributable to OG content. An advantage of SD over LD is its higher hemicellulose content, which can positively impact plant growth by stimulating microbial activity in the rhizosphere, promoting exoenzyme release, and facilitating the mineralization of organic matter ^92,93^. This aligns with evidence suggesting that the type of polysaccharide present in digestate can impact plant growth and productivity by affecting the soil’s physical and biological properties ^94,95^.

Digestate is rich in bacterial and fungal populations, as well as residual plant biomass. Many microorganisms present in digestate thrive under anaerobic conditions but are expected to die when the digestate is stored in open ponds or added to soil. This process can trigger the release of microbe-associated and damage-associated molecular patterns (MAMPs and DAMPs), potentially harming plant growth. Our findings indicate that plants grown in soil treated with microbial-depleted liquid digestate (MD-LD) exhibited improved growth compared to those treated with untreated liquid digestate (LD), while the isolated microbial pellet fraction (M) caused growth inhibition. An advanced DNA metabarcoding approach and taxonomic characterization of the microbial pellet provided deeper insights into its microbial composition and revealed a complex and diverse community structure. Notably, nearly 50% of the sequenced reads were associated with uncultured organisms, highlighting the presence of a substantial fraction of unclassified microbial diversity. Among the predominant genera were *Luteimonas*, *Planomicrobium*, *Caldicoprobacter*, *Pseudomonas*, and *HN-HF0106*. The *Luteimonas* genus, the most abundant bacterial genus in lignocellulosic digestates ^97^, includes species associated with plant growth promotion and pathogen resistance ^98^. *Caldicoprobacter* is highly abundant in the anaerobic digestion of nitrogen-rich substrates ^99^ and is among the most prevalent organisms in 80 digesters ^100^. It is positively associated with *HN-HF0106*, involved in lignocellulose degradation, H₂ and acetate production, and correlated with increased volatile fatty acids in dry anaerobic digestion of corn straw ^101–103^. The first dominant family comprised in the “uncultured” was *DTU014*, a key group involved in biological H₂S oxidation within the digester headspace together with *Caldicoprobacter* and *Syntrophomonadaceae* that were also detected in the samples ^104^. The second most abundant family within the “uncultured” was *Chloroflexi*, a specific native soil microbe ^105^. Its presence in anaerobic digesters was attributed to the persistence of their DNA after their cell lysis ^106^. Up to 15% of uncultured was represented by genera related to *Firmicutes*, a diverse group of bacteria commonly found in soils. As already described in the literature, a large abundance of unaffiliated bacteria genera belonging to the *Firmicutes* are observed in digestate samples ^107^. We found that the olive pomace digestate hosts a highly diverse fungal community. The challenges in assigning fungal communities to taxonomic categories highlight the limitations caused by low DNA concentrations. Similar findings were reported by Langer et al., where the majority of fungal Operational Taxonomic Units remained unclassified in two out of nine digesters analyzed ^108^. A notable number of fungi genetic sequences belonging to *Pichia* and *Basidiomycota* were detected. This result aligns with previous research reporting that the family *Pichiaceae* represented 57% of the *Ascomycota* community in olive mill waste sludge ^109^. Similarly, the 7% of reads mapping to the *Basidiomycota* reference is consistent with findings by Martinez-Gallardo et al., who identified *Basidiomycota* as representing 20% of the microbial community in olive mill wastewater ^110^. The presence of species from the genus *Brettanomyces* was also detected, potentially attributed to their high tolerance to acetic acid ^111^. Of particular interest was the occurrence of genera *Paraglomerales* and *Glomus*, which belong to the arbuscular mycorrhizal fungi, recognized for enhancing plant growth and health by activating defense mechanisms against soilborne pathogens such as *Phytophthora*, *Fusarium*, and *Verticillium* ^112^. The presence of these fungi suggests their resilience during anaerobic digestion, possibly through resistant spores. Fungi could originate from soil residues or plant roots in the raw material. Additionally, post-treatment exposure may have contributed to the colonization by environmental fungi, further increasing the microbial diversity within the digestate. These findings underscore the complexity of the microbial ecosystem present in the digestate, which could influence its biostimulatory properties and agronomic potential. Not all microorganisms in digestate are strict anaerobes; some may survive under aerobic conditions in the soil. The death of bacteria obligate anaerobes—likely contributing to growth inhibition by releasing MAMPs/DAMPs— could be counterbalanced by the survival and activity of aerobic bacteria, which may have a more beneficial effect. A promising approach would be to separate the microbial fractions in digestate, identifying those that enhance plant growth and those that have inhibitory effects. Further analysis is required to identify the key microbes essential for plant development and to determine optimal processing methods for maximizing beneficial effects while minimizing adverse impacts. These findings highlight the necessity of characterizing digestate before agricultural use, ensuring its proper valorization and a comprehensive understanding of the foreign microbial species introduced into the soil.

We hypothesized that microbial molecules and residual plant biomass in digestate could be upcycled to produce bioactive compounds potentially useful as bioelicitors for plant protection against microbial pathogens. A proteinaceous mixture, called MIPE, comprising proteins and peptides from bacterial, fungal, and plant sources, was isolated. Proteins belong to several functional categories, particularly those related to biological processes involved in host interactions and defense responses, suggesting their potential role in enhancing plant immunity. Notably, MIPE contains bacterial and fungal MAMPs such as elongation factor Tu, flagellin, endo-1,4-beta-xylanase A pectate lyase, and histidine kinase. These proteins possess conserved molecular patterns recognized by surface receptors, triggering defense responses that confer broad-spectrum resistance to various pathogens ^113^. For example, application of histidine inhibits wilt disease in tomato and Arabidopsis plants caused by *Ralstonia solanacearum* ^66^. Interestingly, immunogenic peptides like flg22 and elf18 are buried within folded precursor proteins. MIPE included also the plant phytocytokine GOLVEN 1-2A, synthesized as inactive precursors ^68^. Upon MIPE application, plant apoplastic proteases may process these precursors, releasing peptide elicitors recognized as MAMPs by surface receptors, thereby activating immune defenses ^38,114^. The proteomic analysis identifies also a set of CW-degrading enzymes, able to release DAMPs during host-pathogen interaction ^115,116^. The causal agent of citrus canker *Xanthomonas citri* pv. *Citri* synthetizes xylanases generating a broad distribution of xyloglucan oligosaccharides ^69,117^. Pectate lyases can generate OG that are sensed by specific receptors to trigger defense signaling ^118,119^.

Our findings indicate that MIPE acts as a mix of elicitors initiating PTI. MIPE application to Arabidopsis triggered early immune responses including H_2_O_2_ production, MAPK phosphorylation, and upregulation of PTI-related genes. Our evidence indicates that MIPE can trigger PTI signalling pathways similar to those triggered by canonical elicitors, such as flg22 and elf-18 ^31,63,120^. Interestingly, some responses triggered by MIPE, such as FRK1 expression, MAPK4/11 phosphorylation, and H_2_O_2_ production kinetic, appear to be influenced by the presence of multiple elicitors. The synergistic action of different elicitors in MIPE could lead to a broader coverage of immune responses and increased immune efficiency. Indeed, PTI elicitation by flg22 is enhanced when Arabidopsis plants are pretreated with phytocytokine GOLVEN 2 before flg22 application ^68^. The contribution of DAMPs produced by CW-degrading enzymes cannot be excluded and needs further investigation. Several proteomics-identified proteins with unknown functions require further investigation to assess their role in plant immunity. This could reveal new bioactive compounds for plant protection, supporting sustainable agriculture.

Pretreatment of adult Arabidopsis with MIPE primed the expression of defense genes and *CYP81F2*, *PAD3*, and *WRKY53* genes and increased plant protection reducing the lesion area in leaves infected with *B. cinerea* and bacterial growth of *P. syringae*. These findings highlight the potential of MIPE as a universal elicitor of plant immunity, effective not only in model species but also in economically important crops. The presence of a pool of MAMPs, DAMPs, and enzymes releasing DAMPs in MIPE could promote the use of MIPE as plant vaccines to protect plants from diseases. Training plants stress memory is a promising approach to protecting crop reproduction and developing climate-resilient crops. The EU recommends gradually replacing chemical plant protection agents with natural alternatives ^121^. Integrating disease management through pretreatment with natural green elicitors, such as oligosaccharides, is both innovative and sustainable. IBISCO, a chito-oligosaccharide and OG complex, is already on the market, inducing resistance against various pathogens in crops ^122–124^ and is considered a low-risk active substance by the European Commission ^125,126^.

In conclusion, this study highlights for the first time the potential of two-phase olive pomace digestate for sustainable agriculture. It provides a safer alternative to mixed organic digestates, reducing zoonotic pathogen risks. Using two-phase olive pomace as a single biomass ensures composition consistency and a stable microbial community, enhancing protein extraction for plant defense. The solid and liquid fractions can be used as soil conditioners and fertilizer, while the microbial community offers elicitors to boost plant immunity (Figure 9). This valorization process could be applied to digestate wastes from various biomasses, supporting circular economy principles. Future research should optimize biorefining, expand MIPE application to diverse crops, and assess its impact through long-term trials. Regulatory and safety evaluations will be crucial for its adoption, and combining MIPE with sustainable practices like integrated pest management could improve farming solutions.

**Figure 9.**
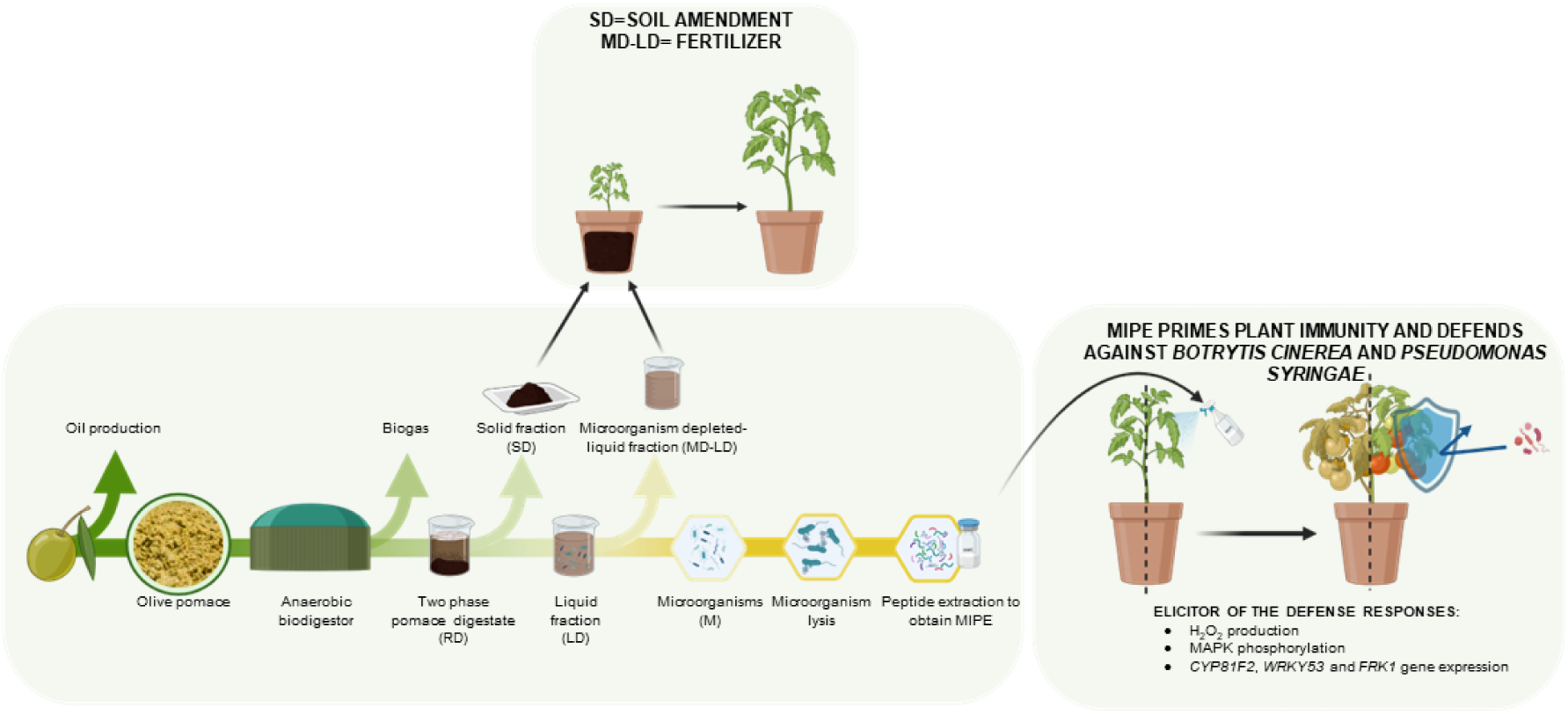
Valorization of digestate for the production of plant defense elicitors. The scheme outlines the valorization process of digestate, a by-product generated during the anaerobic digestion of two-phase olive pomace. Digestate is fractionated into two components: a solid digestate (SD) and a liquid digestate (LD). LD, inherently rich in microorganisms, is subjected to centrifugation to separate the microbial-depleted liquid digestate (MD-LD) from the microbial pellet (M). MD-LD and SD are repurposed as a fertilizer and soil amendment, respectively, enhancing soil fertility and structural integrity. The M undergoes sonication to disrupt the microbial cells, enabling the release of bioactive proteins. The proteins are subsequently purified to produce MIPE, an elicitor capable of priming plant immune responses. Upon application, MIPE triggers key defense mechanisms in plants, including hydrogen peroxide (H₂O₂) production, MAPK phosphorylation, and the upregulation of defense-related genes such as *CYP81F2*, *WRKY53*, and *FRK1*. This strategy has demonstrated efficacy in safeguarding crops against major pathogens, including *Botrytis cinerea* and *Pseudomonas syringae*. By providing a bio-based alternative to synthetic pesticides, this approach underscores the potential of circular bioeconomy strategies in sustainable agriculture.

## AUTHOR CONTRIBUTIONS

**Marco Greco:** Investigation, Methodology, Data curation, Formal analysis, Writing - Original Draft, Visualization. **Daniele Coculo:** Investigation, Writing - Review & Editing, Visualization. **Angela Conti**: Investigation, Writing - Review & Editing, Visualization. **Marco Abatematteo:** Visualization. **Daniela Pontiggia:** Investigation, Writing - Review & Editing, Visualization. **Savino Agresti:** Conceptualization, Resources. **Hugo Mélida**: Writing - Review & Editing, Supervision, Funding acquisition. **Lorenzo Favaro**: Writing - Review & Editing, Supervision. **Vincenzo Lionetti:** Conceptualization, Data curation, Funding acquisition, Investigation, Methodology, Project administration, Resources, Supervision, Validation, Visualization, Writing - Original Draft, Writing - Review & Editing.

## ACKNOWLEDGMENTS

We thank Nicola D’avanzo, Mirko Agresti, Giulia Caminada, and Eleonora Iamele for their technical assistance.

## Supplemental material

**Supplemental Figure S1.**
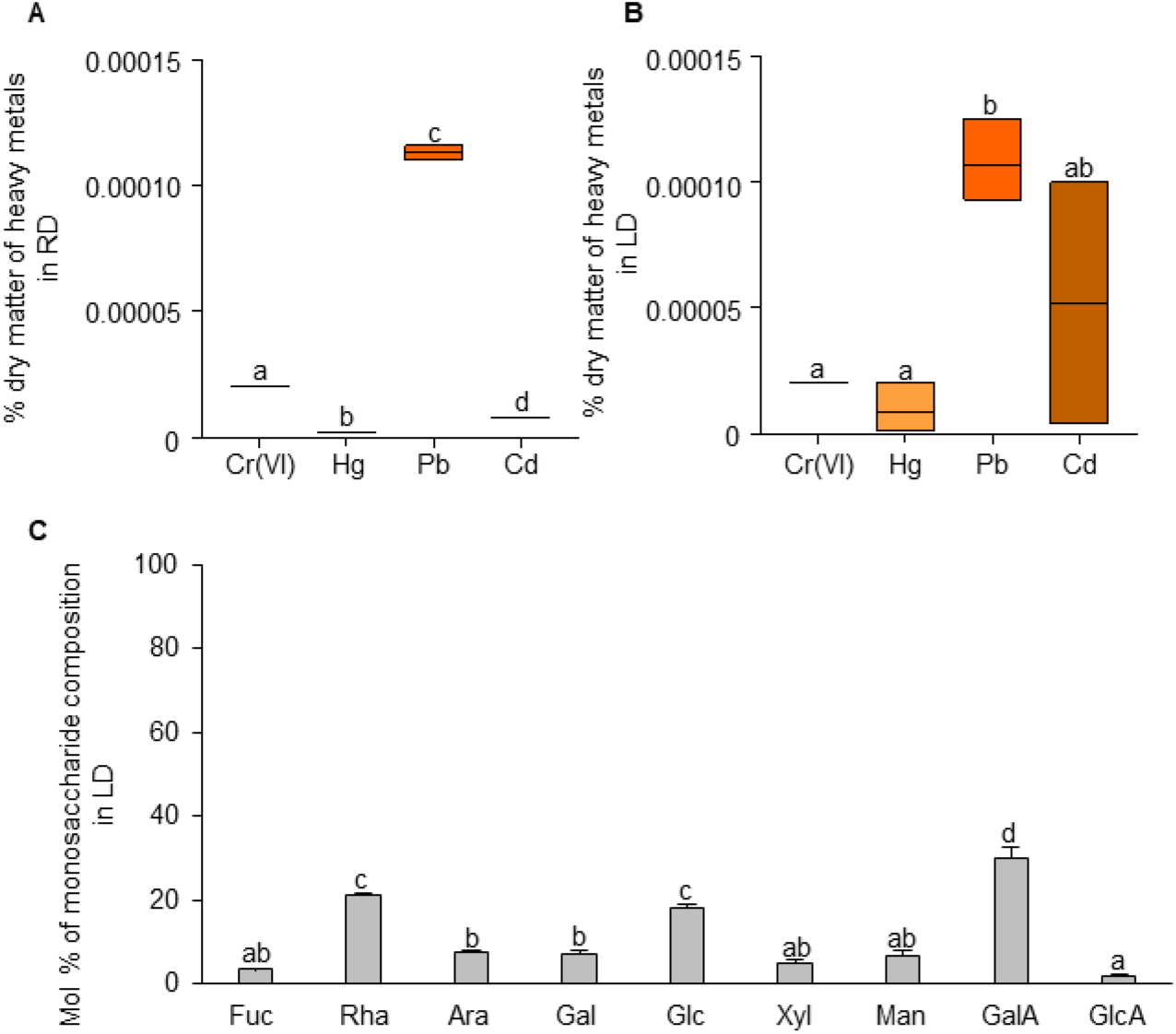
Biochemical composition of RD and LD. (**A**-**B**) Distribution of heavy metals in RD and LD, respectively. Data represent the mean ± standard deviation (n ≥ 3). (**C**) Monosaccharide composition of LD expressed as mol %. Data shown represent the mean ±SD (n=3). The experiments were repeated three times with similar results. The different letters indicate significantly different datasets according to ANOVA followed by Tukey’s test (*p* < 0.05). Chromium hexavalent [Cr(VI)], Mercury (Hg), Lead (Pb), Cadmium (Cd).

**Supplemental Figure S2.**
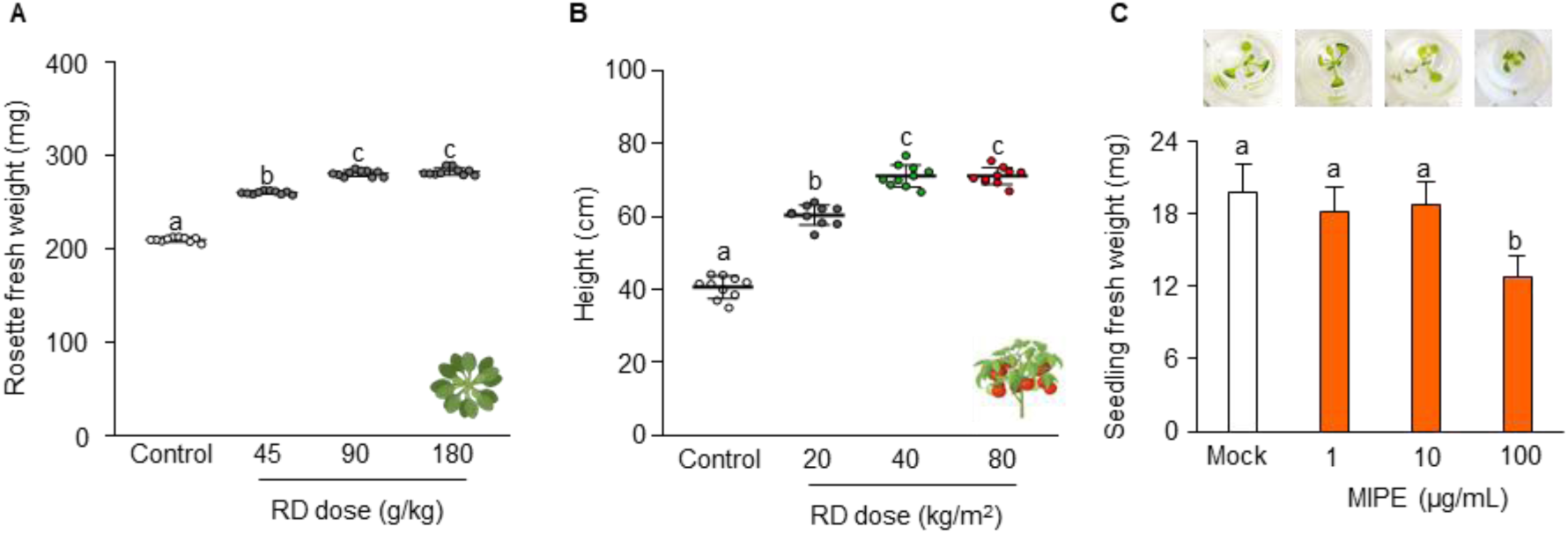
Dose-dependent effects of RD on the growth performance of adult Arabidopsis and tomato, and of MIPE on Arabidopsis seedling growth. (**A**) Dose-effects of soil amended with RD at different doses (45, 90, or 180 g/kg) on *A. thaliana* grown under controlled growth conditions. (**B**) Dose-effects of soil amended with RD (20, 40, or 80 kg/m^2^) on tomato grown on field. (**C**) Effects of water (mock) or MIPE on 10-days-old *A. thaliana* Col-0 seedlings growth. Data shown represent the mean ±SD (n=10). The experiments were repeated three times with similar results. The different letters on the bars indicate significantly different datasets according to ANOVA followed by Tukey’s test (p ≤ 0.05).

**Supplemental Figure S3.**
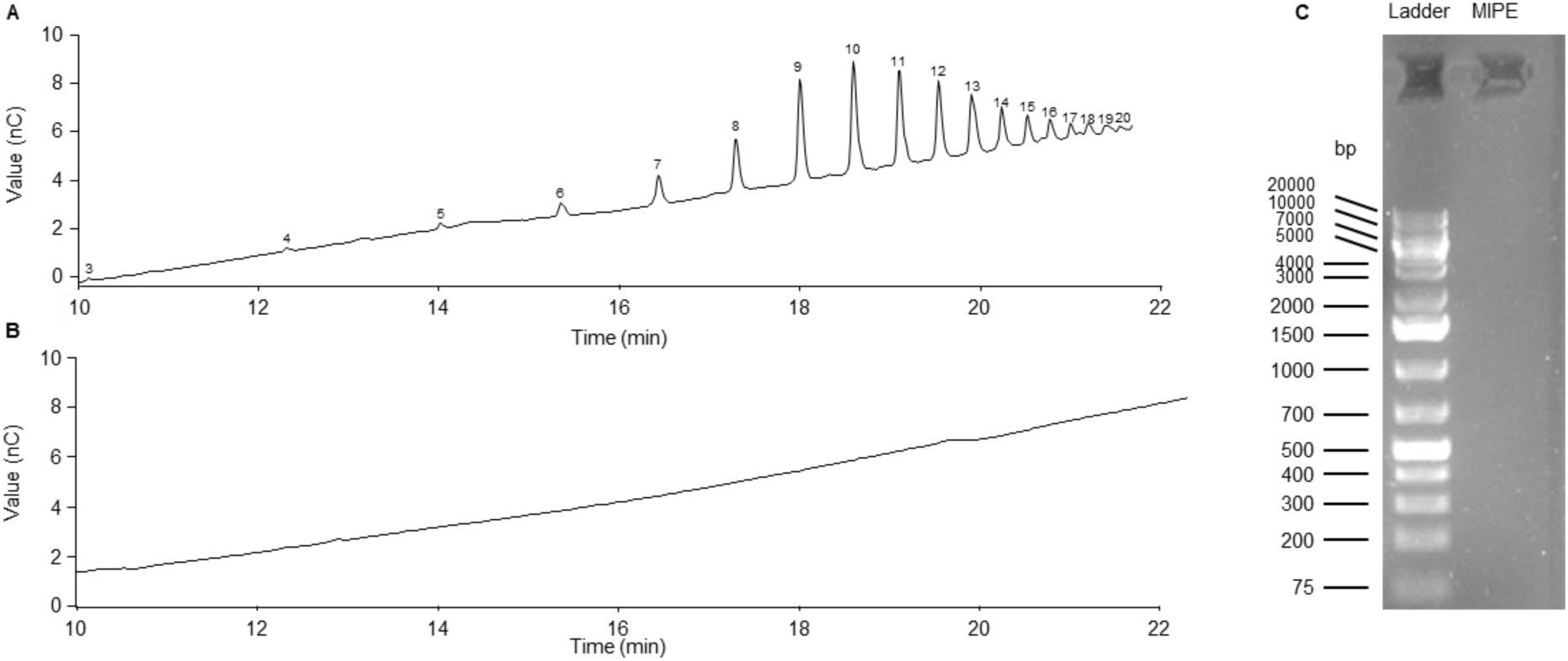
LD and MIPE do not contain OG and nucleic acids, respectively. **(A-B)** HPAEC-PAD profiles of OG used as standard and LD, respectively. The chromatogram shows the intensity of the signals (nC) plotted against retention time (min). Numbers indicate the DP of each OG peak.(**C**) Electrophoresis on a 1.5% (w/v) agarose gel carried out at 60 V for 25 min. Images were captured using Gel Doc™ XR + System (BioRad). Bands indicate nucleic acids at different lengths measured in base pairs (bp).

## Supplemental table

**Table S1.**
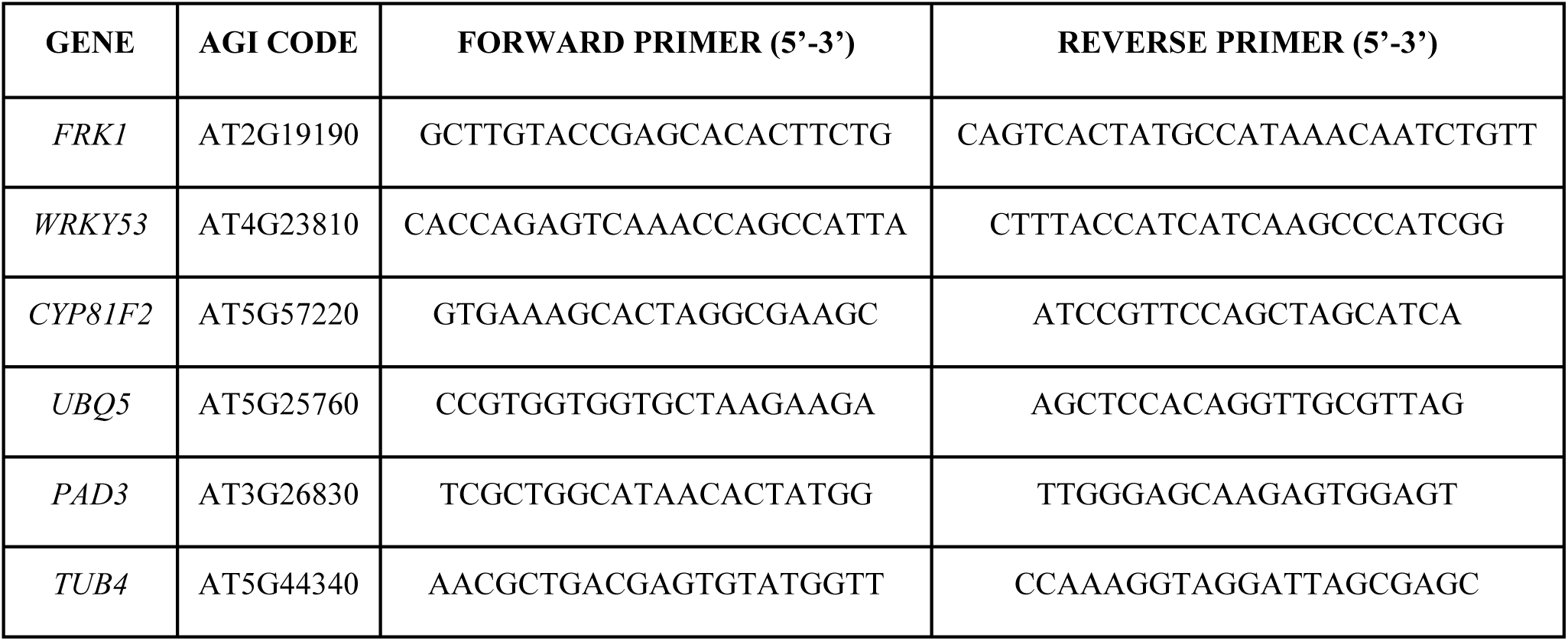
Primers used for Quantitative Reverse Transcript PCR.

